# Beyond coral-algal regimes: high taxonomic resolution surveys and trait-based analyses reveal multiple benthic regimes

**DOI:** 10.1101/2021.01.08.425940

**Authors:** Miriam Reverter, Matthew Jackson, Sven Rohde, Mareen Moeller, Robert Bara, Markus T. Lasut, Marco Segre Reinach, Peter J. Schupp

**Affiliations:** Institute for Chemistry and Biology of the Marine Environment (ICBM) at the University of Oldenburg, Wilhelmshaven, Germany; Faculty of Fisheries and Marine Science, Sam Ratulangi University, Jl. Kampus UNSRAT Bahu, 95115 Manado, Sulawesi Utara, Indonesia; Coral Eye, Bangka Island, North Sulawesi, Indonesia; Helmholtz Institute for Functional Marine Biodiversity at the University of Oldenburg (HIFMB), D-26129 Oldenburg, Germany

**Keywords:** community structure, coral reef regimes, ecosystem functions, functional diversity, traitbased approach, benthic organisms

## Abstract

As coral reef communities change and reorganise in response to anthropogenic and climate disturbances, there is a growing need of detecting and understanding the different emerging species regimes and their contribution to key ecosystem processes. Using a case study on coral reefs at the epicentre of tropical marine biodiversity (North Sulawesi), we explored how application of different biodiversity approaches (i.e. use of major taxonomic categories, high taxonomic resolution categories and trait-based approaches) affects the detection of distinct fish and benthic community assemblages. Our results show that using major categories (family level or above) to study coral reef communities fails to identify distinct regimes. We also show that for detection of different benthic regimes, especially communities dominated by non-coral organisms, monitoring of only scleractinian coral communities is insufficient, and that all types of benthic organisms (e.g. sponges, ascidians, soft corals, algae etc.) need to be considered. We have implemented for the first time, the use of a trait-based approach to study the functional diversity of whole coral reef benthic assemblages, which allowed us to detect five different community regimes, only one of which was dominated by scleractinian corals. We circumvented the challenge that for some benthic groups (e.g. sponges, ascidians or some soft corals) visual identification up to the species level is not possible, by identifying and categorising traits that can be applied to groups of similar organisms instead of specific species. Furthermore, by the parallel study of benthic and fish communities we provide new insights into key processes and functions that might dominate or be compromised in the different community regimes.

## 1. Introduction

Ecosystems worldwide are experiencing profound ecological changes including biodiversity losses (Cardinale et al. 2012) and community rearrangements (i.e. non-random species turnover; Hughes et al. 2018), which are expected to worsen with climate change, even under moderate CO2 mitigation scenarios (Freeman et al. 2013; Guiot and Cramer, 2016). Non-random species turnover, which depends on the susceptibility of the organisms’ traits, can disrupt vital ecosystem processes such as trophic energy flow (Salo et al. 2020) or habitat provisioning (Alvarez-Filip et al. 2011), deeply affecting ecosystem functioning and resilience (Clavel et al. 2011). Good understanding of distinct and emerging species configurations and their contribution to key ecosystem functions is therefore needed to establish effective conservation and management strategies (Darling et al. 2019; Richardson et al. 2020).

Over the past four decades, tropical coral reefs, one of Earth’s most biodiverse ecosystems, have experienced global declines and shifts in species compositions that deeply affect their functioning and the ecosystem services provided (Ainsworth and Mumby, 2015; Hughes et al. 2018; McWilliam et al. 2020). A turnover between highly three-dimensional scleractinian corals such as Acroporidae, by more robust corals (e.g. Poritidae), has been observed worldwide after acute disturbances such as bleaching events or crown-of-thorns outbreaks (Adjeroud et al. 2018; Hughes et al. 2018; McClanahan 2020). Shifts in species compositions including decreases in scleractinians and increases in non-reef building species such as algae, sponges and octocorals are also becoming more frequent as a result of continuous anthropogenic and climate stressors (Bozec and Mumby, 2015; Chaves-Fonnegra et al. 2018; Lasker et al. 2020). Such compositional changes affect several core ecosystem processes (i.e. carbonate production, primary production, trophic interactions and reef replenishment) posing new conservation challenges (Alvarez-Filip et al. 2011; Dixson et al. 2014; Rogers et al. 2014; Brandl et al. 2019).

Coral reefs are extremely heterogeneous ecosystems, with highly varied biological communities that depend on both the local physical environment (e.g. reef topography, wave exposure) and larger biogeographic patterns (Ampou et al. 2017, di Martino et al. 2018). Compositionally and functionally distinct ecosystems will likely respond differently to disturbances, which can then result in different species configurations, further hindering the study and prediction of coral reef trajectories and their effect on core ecosystem processes (Roff and Mumby 2012; Ampou et al. 2017). In this context, conservation approaches need to consider both coral reefs spatio-temporal heterogeneity (i.e. different species configurations) and their contribution towards core ecosystem processes.

The study of species configurations (i.e. community biodiversity) has been traditionally studied as the relative abundance of different species, and as such, most studies assessing coral reef composition have mostly used taxonomic categories (often at family level or higher, especially for benthic organisms) to identify community changes (e.g. Jouffray et al. 2015; Reverter et al. 2020). Approaches using major taxonomic categories (e.g. hard coral, soft coral, algae, etc.), have the advantage of being easily implemented in global citizen science programs such as ReefCheck (Schläppy et al. 2017; Vallès et al. 2019), and have allowed identification of marked regime shifts, for example, from coral to algae-dominated communities (e.g. Bakker et al. 2017; Reverter et al. 2020). However, the use of major benthic categories might overlook functionally important compositional changes (McClanahan 2017; Gonzalez-Barrios et al. 2020). Many studies have shown that different species contribute differently to ecosystem functioning, and therefore ecological research has seen a shift from taxonomic diversity to functional diversity studies (Darling et al. 2012; Mouillot et al. 2013). Trait-based approaches offer new opportunities for deeper mechanistic understanding on the role of biodiversity in maintaining multiple ecosystem processes and they allow identification of species with critical and vulnerable ecosystem functions (Mouillot et al. 2014; Madin et al. 2016). Trait-based approaches have been successfully used to study the changes in coral reef fish communities (e.g. Mouillot et al. 2014; Ottimofiore et al. 2017; Richardson et al. 2018) and in scleractinian coral assemblages (e.g. Darling et al. 2012; Denis et al. 2017; Kubicek et al. 2019). However, whilst scleractinian corals are the key organisms of coral reefs, recent shifts towards assemblages dominated by alternate organisms, highlights the need of expanding these approaches to include all types of benthic organisms.

Here, we studied coral reefs around Bangka and Bunaken islands (North Sulawesi, Indonesia), which are at the epicentre of marine biodiversity (Hoeksema 2007) and display high spatial heterogeneity (Ponti et al. 2016; Ampou et al. 2017). We used them as a case study to explore how the use of different biodiversity approaches (major taxonomic categories, high taxonomic resolution categories and trait-based approaches) affects the detection of distinct community (benthic and fish) compositions. We also implemented, for the first time, the use of a trait-based approach to study the functional diversity of coral reef benthic assemblages, including all types of sessile organisms encountered (e.g. sponges, ascidians, soft corals, etc.). We also analysed how the determination of benthic communities and the identification of distinct configurations change when commonly used groups such as scleractinian corals or reef fish are complemented with all sessile benthic organisms.

## 2. Material and methods

### Coral reef surveys

Benthic and fish surveys were conducted at nine different sites at Bunaken and Bangka islands (North Sulawesi, Indonesia) between February and March 2020 (Figure 1). The sites were chosen due to their high heterogeneity (i.e. reef topography, exposure and anthropogenic impacts), in order to reflect different community regimes around these two islands (Supplementary Table 1). Three 20-m transects (separated by 5 m) were placed at each site parallel to the coast at either 3 or 10 m deep (Figure 1). The benthos and fish communities were characterised on each transect using three different levels of ecological information: major categories, which have been used in citizen science programs such as ReefCheck, categories at the highest taxonomic resolution possible (e.g. genus or species) and functional entities (Supplementary methods).

**Figure 1.**
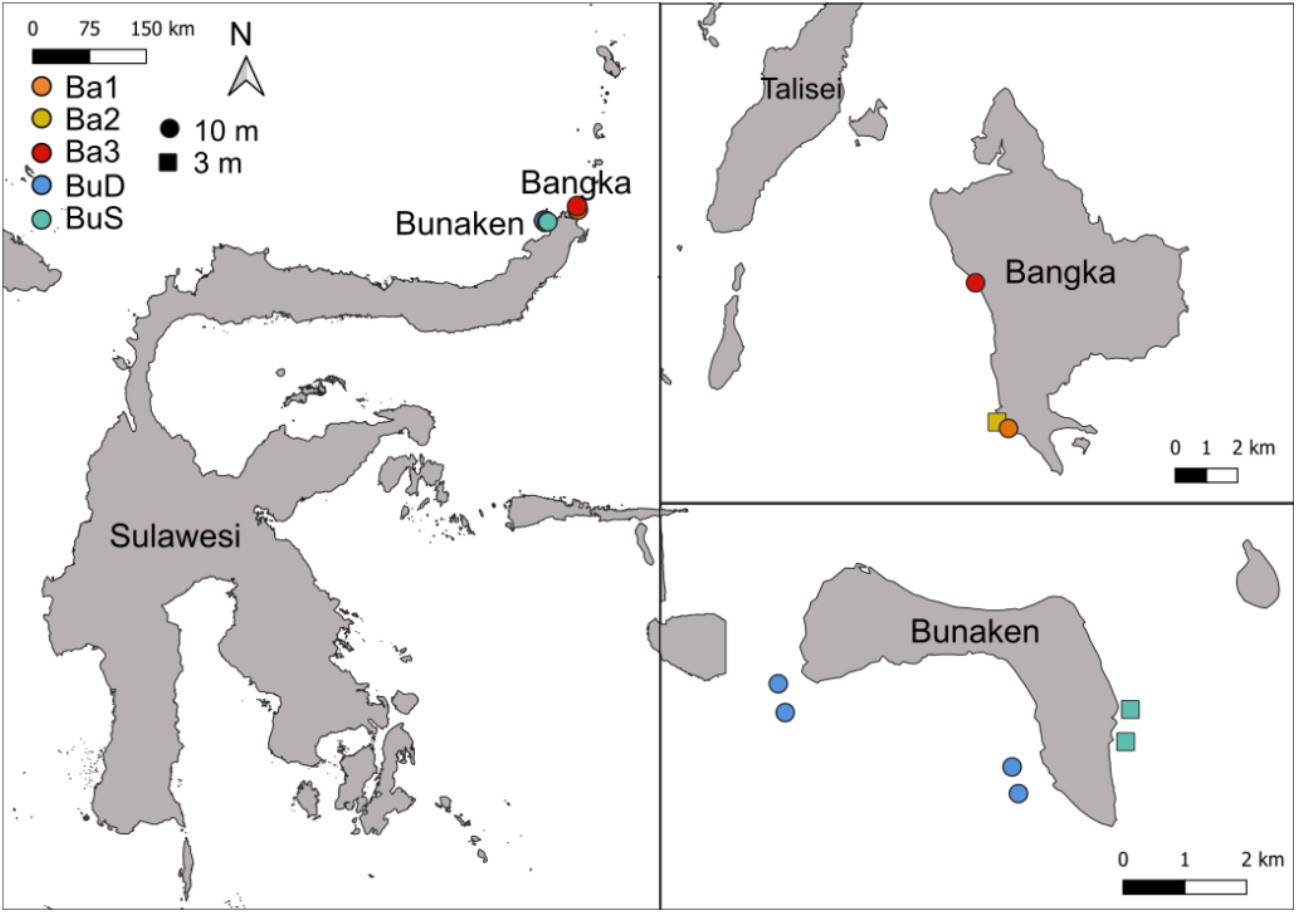
Map of the sites monitored around Bangka and Bunaken Islands (North Sulawesi, Indonesia). Colours indicate the different regimes identified.

For the benthos composition study, we photographed the reef bottom every 0.5 m from a distance of 0.5 m above the benthos using a camera (Olympus TG-5) mounted on a squared metal frame with a surface area of 0.25 m^2^. All photographs were analysed using the software Coral Point Count Excel extension (Kohler and Gill 2006) to determine the percentage cover of different sessile organisms. Thirty random points were assigned to each photograph and the organisms under these points were identified to the lowest taxonomic level possible (n = 120 points per transect, 99 benthic categories identified, Supplementary table 2). These taxonomic categories were classified into seven major categories: hard coral, soft coral, sponge, coralline algae, macroalgae, fleshy algae and other living organisms. Scleractinian corals were also classified into 11 sub-categories according to their morphology: branching, caespitose, columnar, corymbose, digitate, foliose, massive, submassive, solitary and tabular, which were used as major categories in the analyses of only scleractinian coral communities (Supplementary table 2, Supplementary methods).

For the fish survey, 3-min videos were taken swimming along the transects at a constant speed. A first video was filmed while laying the transect and was used to collect data on bigger fish species that might be scared away by the divers. A second fish video was filmed five minutes after laying the transect and was used to count the resident fish species. All fishes (> 5 cm) within 3 m on each side of the transect and 3 m above were counted, their family and species identified and their approximate length (± 3 cm) recorded. Larger and rarer fish (e.g. unicornfishes, parrotfishes, big groupers, pelagic fish such as barracudas and trevallies) were counted when observed within 5 m from the transect. Fish biomass was calculated following Friedlander & DeMartini (2002) using the equation W = a x L^b^, where W is the weight of the fish in grams, L is the total length (LT) in cm and a and b are species-specific constants obtained from FishBase (Froese & Pauly, 2014). Fish families were used as major categories, whereas fish species were used for the high taxonomic resolution analysis.

### Benthos and fish traits

The functional ecology of benthos categories was characterised using 12 traits: colony formation, growth form, maximum colony size, longevity, growth rate, body flexibility, skeleton presence, reproductive strategy, sexual system, feeding strategy, presence of photosynthetic symbionts and corallite maximum width (only for scleractinian corals). The chosen functional traits focus on key ecosystem processes that affect the organisms’ population dynamics, coral reef accretion and nutrient cycling and resources. Since category identification was performed at genus or higher taxonomic ranks, we often used ordered categories to classify the quantitative traits (Supplementary Methods). Most of the functional traits from scleractinian corals were extracted from the Coral Trait Database (Madin et al. 2016; https://coraltraits.org/), whereas scientific references and monographs were used to extract information on the other organisms traits (Supplementary Table 2).

The functional ecology of fish species was characterised using six traits (body size, diet, period of activity, vertical position, gregariousness and mobility) (Supplementary Methods, Supplementary Table 3). The chosen traits describe the main facets of fish ecology and are relevant in critical ecosystem processes such as nutrient cycling and food web regulation (Mouillot et al. 2014). Fish trait data was collected from the FishBase database (www.fishbase.org) and from published articles studying coral reef fish functions (Mouillot et al. 2014; MacNeil et al. 2015; Bierwagen et al. 2018). Unique combinations of traits were defined as functional entities (FEs).

### Multivariate community analysis

Benthic cover and fish biomass data were transformed using the Hellinger transformation (function “decostand” from the vegan R package, Oksanen et al. 2019) prior to the multivariate analyses. Nonmetric multidimensional scaling (NMDS, metaMDS function from the vegan package) were used to visualize the differences in benthos (scleractinian corals and all benthic animals) and fish communities between the different sites. The three levels of ecological information (major categories, categories defined at the lowest taxonomic level possible and functional entities) previously described were used to investigate how they affect community pattern detection. A hierarchical cluster analysis (function hclust in R) using Euclidian distance was then used to identify clusters of similar sites. The “Average” algorithm was chosen after analysis of the cophenetic correlation coefficient (Pearson correlation between the cophenetic distances calculated on cluster branches and the benthos/fish community dissimilarity matrix). The Kelley-Gardner-Sutcliffe (KGS) penalty function (maptree R package, White and Gramacy, 2015) was used to prune the dendrogram. The different clusters identified were considered hereafter as compositionally different communities and were used for the subsequent description of the communities using high taxonomic resolution data and functional analysis.

### Functional space and functional indices

The functional richness was calculated as the volume within the multidimensional functional space enclosing all the FE in a specific community, where each species is placed according to their functional niche (Villeger et al. 2008; Maire et al. 2015). First, a species dissimilarity matrix was built using the Gower’s distance (Gower, 1971; Pavoine et al. 2009), which can deal with all types of traits (continuous, ordinal and categorical), and gives them the same weight. Then, a Principal Coordinates Analysis (PCoA) was performed using the previous dissimilarity matrix. In order to select the number of PCoA axis that would result in the best functional space, which needs to be congruent with the initial functional distance, we computed the mean squared deviations (mSD) of functional spaces with multiple axis (up to 10), in which lower mSD represents a higher quality of the functional space (Maire et al. 2015). After examination of mSDs, we selected four axis to build both of our functional spaces (benthos and fish), since adding a fifth axis only weakly increased the quality of the functional spaces (Supplementary figures 2, 3). The functional space and the mSD values were computed using the R function quality_funct_space, developed by Maire et al. (2015).

The number of FE (FErichness) was calculated to explore the functional diversity. We also computed the community-weighted means of trait values (CMW) using the dbFD function from the FD R package (Laliberté et al. 2015). The CMW provides information on functional composition by identifying the most common value traits in a specific community.

## 3. Results

### Identifying community patterns

We studied the benthic (scleractinian corals and all benthic organisms) and fish community composition of nine coral reefs in North Sulawesi (Indonesia) by using major categories, categories at the highest taxonomic resolution possible and FEs (Supplementary table 2, 3). The analysis of high taxonomic data and functional entities showed marked differences between the benthos (all benthic organisms) and fish communities of the sites studied, allowing the identification of five significantly different clusters (i.e. sites with similar compositions) (Figure 1, 2, Supplementary Figure 1). Each of the sites in Bangka island formed its own cluster (Ba1, Ba2 and Ba3), whereas the sites at Bunaken where grouped into two clusters containing the deep (Bu D cluster containing Bu1, Bu2, Bu3 and Bu4) and shallow sites (BuS cluster, containing Bu5 and Bu6) (Figure 1, 2, Supplementary Figure 1).

**Figure 2.**
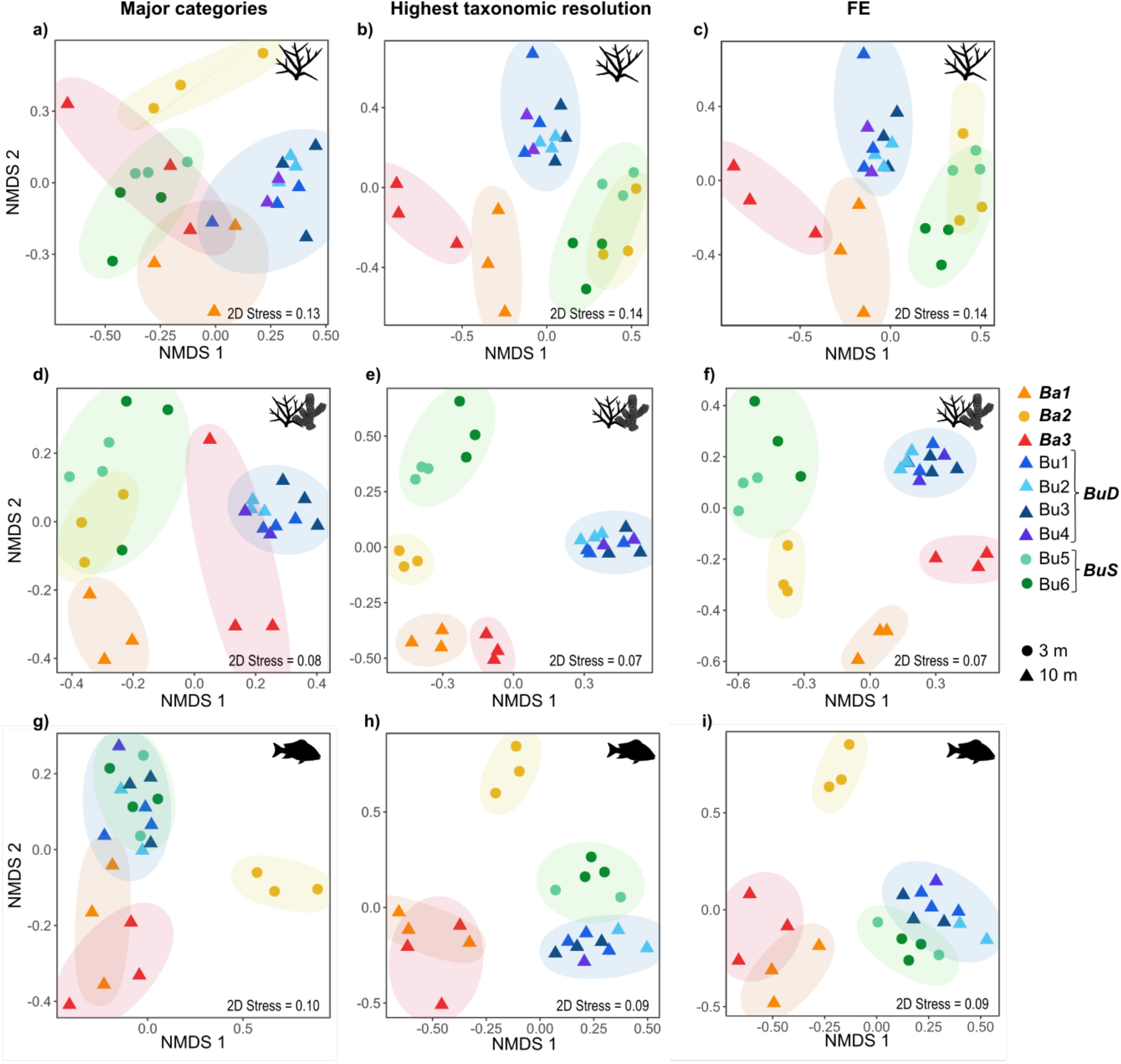
Analysis of the benthos (only scleractinian corals a-c and all benthic organisms d-f) and fish (g-i) community similarities between the different sites samples using three levels of ecological information: major categories, highest taxonomic resolution and functional entities (FE). Community clusters of sites grouping together are highlighted in bold and italics (Ba1, Ba2, Ba3, BuD and BuS).

Regardless of the type of community studied (i.e. only scleractinian corals, all benthic organisms or fish) the use of major categories failed to identify most of the community differences highlighted by the use of higher resolution data (i.e. high taxonomic resolution or functional entities) (Figure 2). For example, the analysis of coral major categories (i.e. classified upon their morphology), only allowed the identification of BuD community, whereas all the other Bunaken and Bangka sites were mixed (Figure 2a, Supplementary Figure 1). Similarly, the analysis of the coral reef benthos using major categories only identified the BuS and Ba3 clusters and the fish family analysis only allowed the identification of significantly different communities in Ba2 cluster, but grouped together the other Bangka sites (Ba1 and Ba3) and all Bunaken sites (Figure 1, 2, Supplementary Figure 1).

The results also show that the study of all benthic organisms allows much better detection of different community regimes than just the study of scleractinian coral communities. For example, the sites from Ba2 grouped together with the shallow Bunaken sites (BuS) when using only scleractinian corals, but were clearly distinguished when all the benthic organisms were considered (Figure 2, Supplementary Figure 1).

### Benthic community structure

Our dataset was composed of highly heterogeneous reefs, with each of the communities detected dominated by different benthic organisms: scleractinian hard corals (BuS), blue coral *Heliopora coerulea* (Ba2), xenid soft corals (Ba1), colonial ascidians (Ba3) and sponges (BuD) (Figure 3a, b).

**Figure 3.**
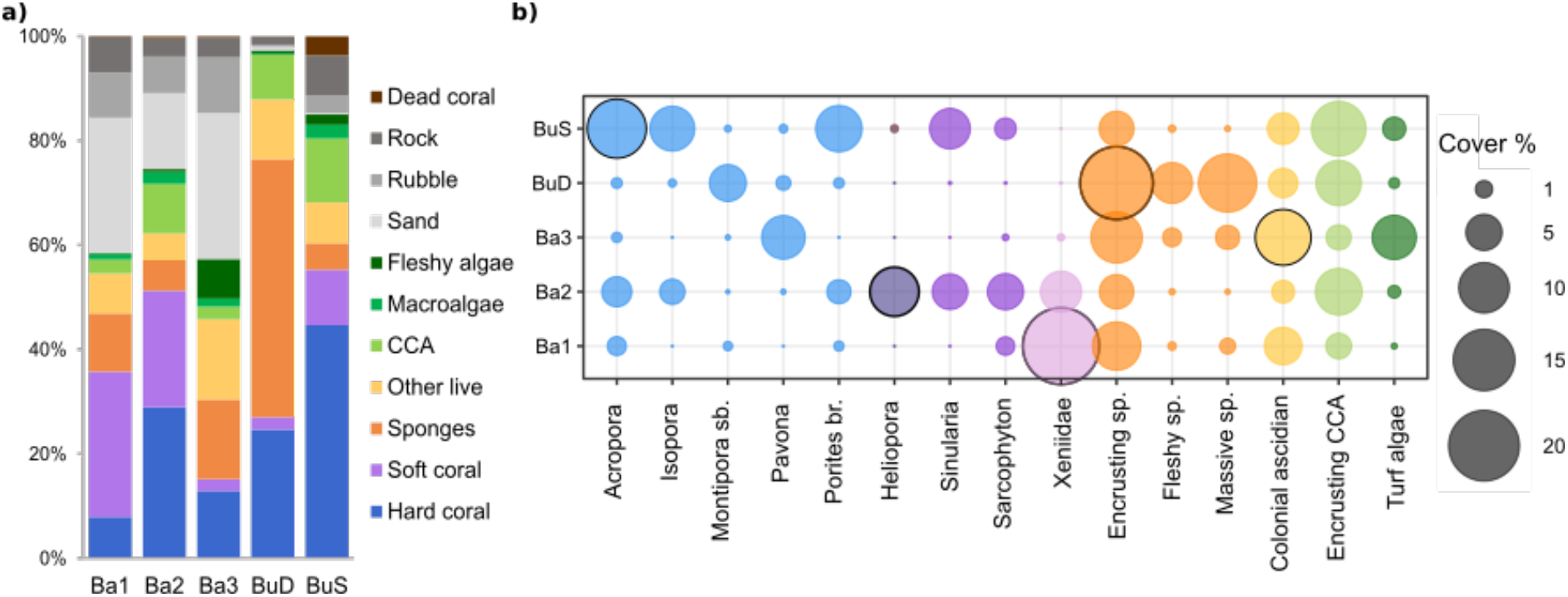
Benthic composition (based on major categories) of the different community clusters identified (a) and cover (%) of the most abundant (≥ 5% at least in one cluster) benthic taxonomic categories identified (b). Highlighted circles represent the dominant benthic organism in each of the communities.

Only two (Ba2 and BuS) out of the five community clusters identified were dominated by reef-building species (i.e. hard corals, 28.9 ± 6.5% and 44.7 ± 15.6%, respectively). The reef at Ba2 was predominantly dominated by the blue coral *H. coerulea* (9.5 ± 3.7%), with lower covers of branching scleractinian corals (*Acropora* spp. 3.4 ± 3.5% and *Porites* spp. 2.1 ± 0.6%) and columnar coral *Isopora palifera* (2.4 ± 2.4%) (Figure 3a, 3b). Ba2 also displayed important covers of encrusting coralline algae (8.5 ± 0.3%) and soft corals from the Alcyoniidae (mainly *Sarcophyton* spp. 5.0 ± 3.4% and *Sinularia* spp. 4.8 ± 2.1%) and Xeniidae (6.6 ± 2.3%) families. BuS reefs were dominated by branching scleractinian corals (*Acropora* spp. 13.7 ± 16.9% and *Porites* spp. 8.1 ± 6%), the columnar coral *Isopora palifera* (7.8 ± 5.9%) and massive *Porites* spp. (4.4 ± 4.0%). BuS sites also displayed large covers of encrusting coralline algae (11.7 ± 2.6%) and Alcyoniidae corals (mostly *Sinularia* spp. 6.3 ± 5.9%) (Figure 3a, 3b).

Ba1 transects were dominated by soft corals from the Xeniidae family (23.5 ± 3.0%), followed by incrusting sponges (9.2 ± 3.6%), and presented low hard coral cover (7.9 ± 1.3%). Ba3 was dominated by sponges (15.2 ± 3.1%), mostly encrusting sponges (10.3 ± 1.3%), followed by ascidians (13.4 ± 4.4%, mostly incrusting colonial ascidians) and hard corals (12.9 ± 8.1%, mostly *Pavona* spp.). Sponges accounted for 49.4 ± 6.5% of the cover in BuD transects, with encrusting sponges being the most abundant organisms (21.2 ± 4.2%), followed by massive (13.4 ± 3.8%) and fleshy incrusting sponges (6.4 ± 1.5%). Hard coral cover was 24.7 ± 4.9%, with submassive *Montipora* ssp. (4.2 ± 2.9) and massive *Porites* spp. (3.1 ± 1.4%) being the most abundant genus (Figure 3a, 3b).

### Functional diversity analysis of benthos communities

The 99 benthic categories were classified into 64 FEs (Supplementary Table 2) for which their functional niche was displayed using a functional space built on four PCoA axis. Generally, species longevity, and corallite maximum width (for scleractinian corals) decreased along the first axis (PC1), whilst flexibility and growth rate tended to increase with PC1 (Figure 4a). Colony form was highly structured along the fourth axis (PC4), with encrusting species at the top, branching species at the middle and massive species at the bottom of PC4 (Figure 4a). 29 out of the 64 FEs contained calcified species contributing to reef accretion (e.g. hard corals, crustose coralline algae, foraminifera), with nine FEs also contributing to reef structural complexity (i.e. branching morphology). 25 FEs out of the 64 FEs identified contained fast-growing species, including some potentially proliferating species (e.g. cyanobacteria, macroalgae, encrusting sponges, encrusting ascidians).

**Figure 4.**
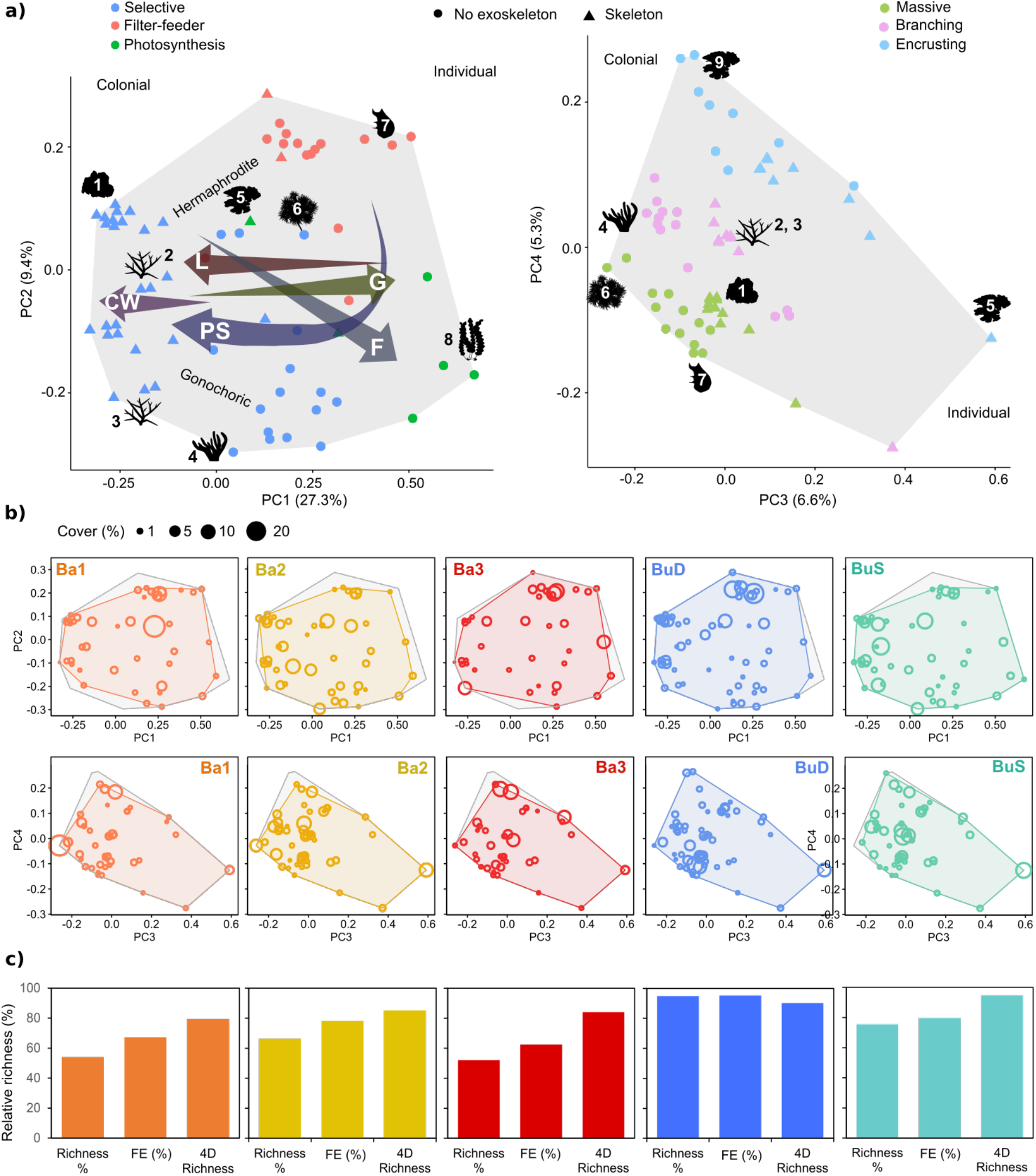
Benthic functional diversity of the different communities identified. a) Distribution of functional entities (FEs) in the global benthic functional space, built using four PCoA axis (PC1 and PC2 left, PC3 and PC4 right) using twelve functional traits: colony formation, growth form, maximum colony size, longevity (L), growth rate (G), body flexibility (F), skeleton presence, reproductive strategy, sexual system, feeding strategy, presence of photosynthetic symbionts (PS) and corallite maximum width (CW, only for scleractinian corals). The numbers indicate the following functional entities: 1: massive hermaphrodite scleractinian corals, 2: branching hermaphrodite scleractinian corals, 3: branching gonochoric scleractinian corals, 4: *Sinularia* soft coral, 5: encrusting crustose coralline algae, 6: Xeniidae soft coral, 7: solitary ascidians, 8: macroalgae, 9: encrusting filter-feeders (sponges and ascidians). b) functional spaces of each of the communities analysed (coloured convex hull) superposed to the global functional space. c) functional diversity indices for each of the communities: relative taxonomic richness (Richness %), relative FE richness (FE %) and relative functional richness as % of filled global functional space (4D richness).

The three Bangka sites displayed the lowest taxonomic and FE richness (S_richness_ = 52-67%, FE_richness_ = 63-78%), but still filled 80 (Ba1), 85 (Ba2) and 84% (Ba3) of the benthos functional space (Figure 4b, c). Only Ba2 was characterised by reef-building species (branching long-lived calcified species with fast growths) (Table 1, Figure 4b). Ba2 contained 24 out of the 29s FEs contributing to reef accretion and the 9 FEs also contributing to reef structural complexity (Figure 4b). Ba1 and Ba3 were in contrast, characterised by fast-growing short-lived species with no contribution to reef accretion (i.e. no skeleton) (Table 1, Figure 4b). Ba1 and Ba3 also displayed the lowest functional diversity of reef-building species, with 19 and 18 FEs contributing to reef accretion, respectively. Ba1 contained only five FEs contributing simultaneously to reef accretion and reef structural complexity, but the cover of these FEs was extremely low (< 3%). Ba3 contained only four branching reef-building FEs, with the FE containing *Pavona* spp. reaching 7% of the benthic cover. Ba3 also displayed the lowest cover in hermaphrodite broadcaster branching hard corals such as Acroporidae (Figure 4b).

**Table 1.** Community-weighted-mean values (CWM) for the different traits and benthic communities studied. FS = feeding strategy, RS = reproductive strategy, CMW = corallite maximum width, GR = growth rate.

| Site cluster | Colonial | Form | Flexibility | FS | Photosynthetic symbionts | RS | Sexual system | Skeleton | CMW | GR | Colony size | Longevity |
| --- | --- | --- | --- | --- | --- | --- | --- | --- | --- | --- | --- | --- |
| Ba1 | Y | Mass. | 3 | Sel. | 3 | Br | H | N | 1 | 3 | 1 | 2 |
| Ba2 | Y | Bra. | 1 | Sel. | 3 | Bc | G | Y | 1 | 3 | 3 | 4 |
| Ba3 | Y | Encr. | 1 | FF | 2 | Al | H | N | 1 | 3 | 2 | 1 |
| BuD | Y | Mass. | 1 | FF | 2 | Al | H | N | 1 | 1 | 3 | 4 |
| BuS | Y | Bra. | 1 | Sel. | 3 | Bc | H | Y | 1 | 1 | 3 | 4 |

Body form categories: Mass. = massive, Bra. = branching, Encr. = encrusting. FS categories: Sel. = selective, FF = filter-feeder. RS categories: Br = brooder, Bc. = broadcasters, Al. = alternate.

BuD presented the highest taxonomic and FE richness (95%), however presented a smaller functional richness (90%) than BuS (95%) (Figure 4b, c). BuD sites were dominated by massive, long-lived, slow-growing filter-feeding species, such as barrel and massive sponges; whilst BuS was characterised by branching, calcified, long-lived, broadcaster species such as Acroporidae corals (Table 1, Figure 4b). Both BuD and BuS contained most of the FEs contributing to reef accretion and reef structural complexity (Figure 4b).

Out of the 64 FEs, only 29 FEs were found in all sites (Figure 5). BuD was the site with the highest number of unique FEs (9), but none of the Bangka sites presented any unique benthic FEs (Figure 5). Bunaken communities (BuD and BuS) presented 2 FEs that were absent from the sites studied at Bangka island (Figure 5). Deep sites (BuS, Ba1 and Ba3) presented two FEs that were absent in the shallow sites, whereas the shallow communities (BuS and Ba2) had one unique FE, the blue coral *H. coerulea* (Figure 5).

**Figure 5.**
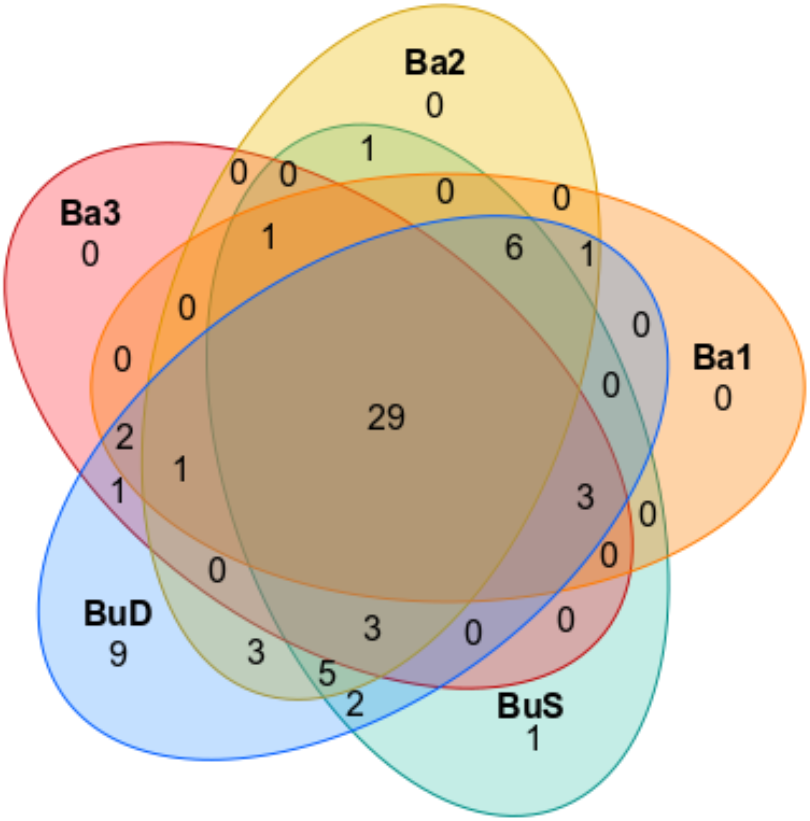
Venn diagram representing the shared functional entities (FEs) between the different communities detected. Ba: Bangka sites, Bu: Bunaken sites.

### Functional diversity analysis of fish communities

The 172 fish species were classified into 97 FEs (Supplementary Table 3), for which their functional niche was displayed using a functional space built on four PCoA axis. Generally, gregariousness and vertical position increased along the first axis of the functional space (PC1) (Figure 6a). The second axis (PC2) was characterised by differences between nocturnal and diurnal species, fish size and diet, showing a clear separation between planktivorous (e.g. damslefishes, fusiliers) and piscivorous fish (e.g. snappers, barracudas). Fish mobility increased along PC1 and decreased with PC2. PC3 showed a clear separation between nocturnal (left) and diurnal (right) species, with mobility generally increasing with PC3 values, with highly mobile fish species such as fusiliers or surgeonfishes at the right extreme of the functional space. Fish gregariousness generally decreased along PC4, but the pattern was not as clear as with PC1. Vertical position also tended to increase with PC4, with highly substrate associated species such as parrotfishes or squirrelfishes at the bottom of the functional space (Figure 6a).

**Figure 6.**
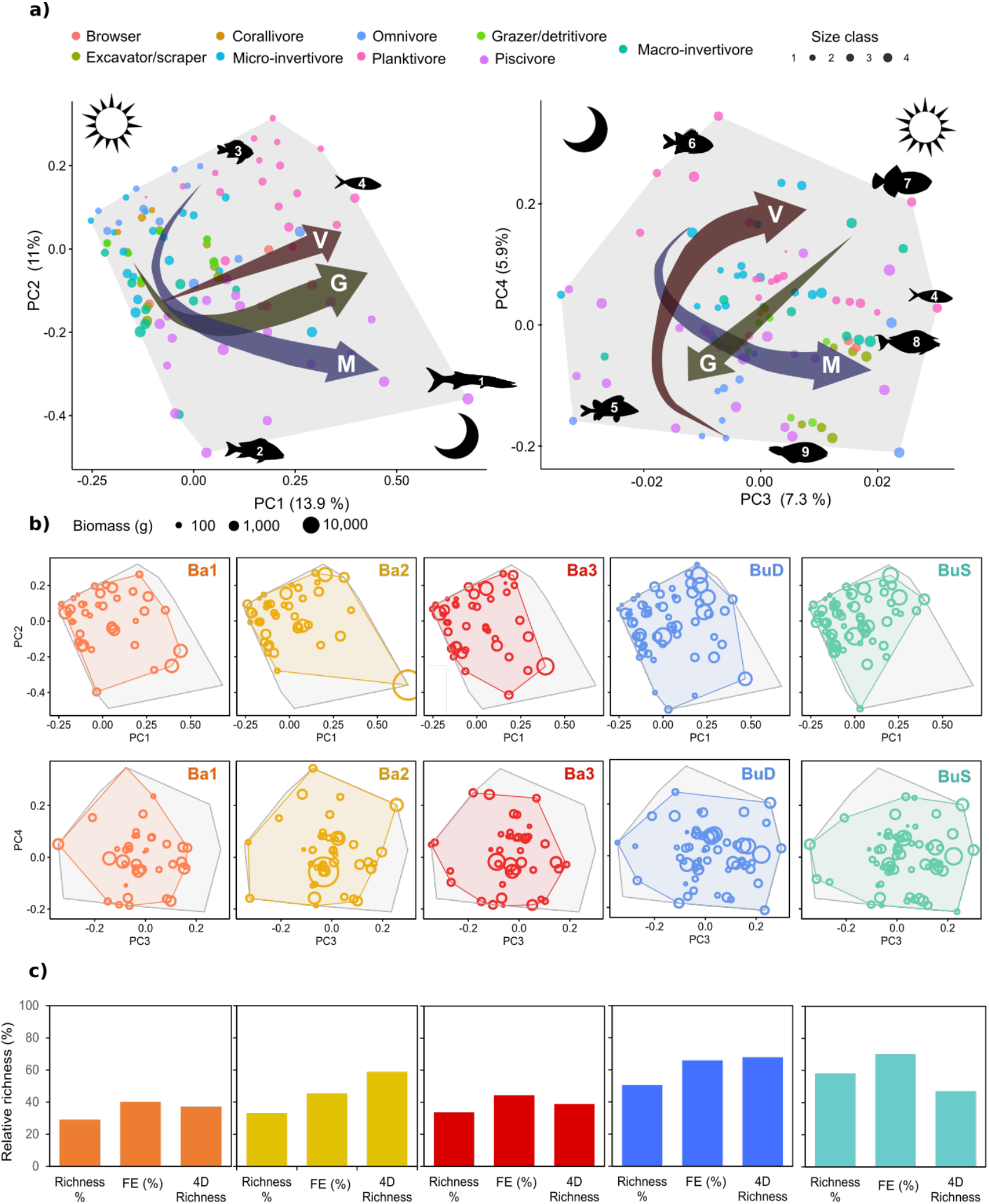
Fish functional diversity of the different communities identified. a) Distribution of functional entities (FEs) in the global fish functional space, built using four PCoA axis (PC1 and PC2 left, PC3 and PC4 right) using six functional traits: body size, diet, period of activity, vertical position (V), gregariousness (G) and mobility (M). The numbers indicate the following functional entities: 1: *Sphyraena quenie* (Sphyraenidae), 2: big snappers (e.g. *Macolor macularis*, Lutjanidae), 3: damselfishes, 4: pelagic planktivores such as fusiliers (Caesionidae), 5: squirrelfishes (*Sargocentron* spp., Holocentridae), 6: soldierfishes (*Myripristis* spp., Holocentridae), 7: *Melichthys vidua*, 8: unicornfishes (*Naso* spp, Acanthuridae) b) Functional spaces of each of the communities’ analysed (coloured convex hull) superposed to the global functional space (grey). The bubble sizes represent the FEs mean biomass at each of the clusters c) Functional diversity indices for each of the communities: relative taxonomic richness (Richness %), relative FE richness (FE %) and relative functional richness as % of filled global functional space (4D richness).

All Bangka sites displayed the lowest taxonomic and FE richness (S_richness_ = 29-33%, FE_richness_ = 4045%). Ba1 and Ba3 also presented the lowest functional richness, filling only 37 and 39% of the functional space respectively. Ba2, however, exhibited the second highest functional richness, filling 59% of the functional space (Figure 6b, c), which was related to the presence of few highly original FEs such as big highly mobile predators (i.e. barracudas, PC1) and the nocturnal, gregarious, highly-site attached omnivorous sweeper (*Pempheris oualensis*, PC3-PC4), which were uniquely found in Ba2 (Figure 6b). Overall, all Bangka sites were characterised by the lack of browsers; although this trophic group was only represented by two FE englobing three Acanthuridae species (Supplementary table 3). Ba1 and Ba3 also displayed low functional diversity and biomass of grazers/detritivores. Ba1 and Ba2 lacked the presence of most big high-trophic chain fish (i.e. piscivorous, macro-invertivore and omnivorous), excepting the barracudas in Ba2 (Figure 6c).

Bunaken sites harboured the highest number of fish species and FEs, which were characterised by middle-size, diurnal planktonic species (Table 1, Figure 6b). BuD, which harboured 50% of all species and 66% of all FEs, displayed the highest functional richness (68%). BuS, which hosted the largest taxonomic richness (58%) and FE richness (70%) only filled 47% of the functional space, which was related to the absence of nocturnal piscivorous species such as snappers or barracudas (Figure 6c).

Out of the 97 FEs identified in all communities, only 19 FEs were present in all sites (Figure 7). None of the piscivore FEs was shared between all sites, but, all sites presented unique piscivore FEs. BuD was the community with the highest number of unique FEs (11), followed by BuS (9) and then the Bangka sites Ba3 (4), Ba2 (3) and Ba1 (3) (Figure 7). Bunaken communities had 9 FEs that were absent in the communities studied in Bangka island, which were mostly medium to highly mobile FEs such as fusiliers (Caesionidae) and large Acanthuridae (Figure 6, 7). Shallow sites (BuS and Ba2) presented four unique FEs that were absent from deep sites (BuS, Ba1 and Ba3) (Figure 7).

**Figure 7.**
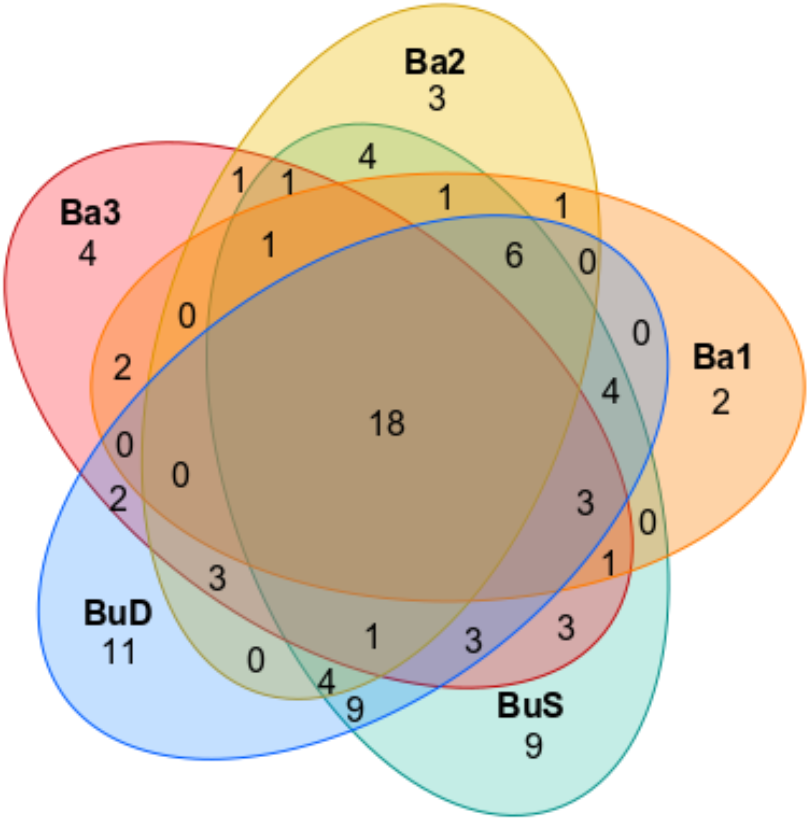
Venn diagram representing the shared functional entities (FEs) between the different communities detected. Ba: Bangka sites, Bu: Bunaken sites.

**Table 2.** Community-weighted-mean values (CWM) for the different traits and fish communities studied.

| Site cluster | Size | Mobility | Activity | Gregariousness | Vertical position | Diet |
| --- | --- | --- | --- | --- | --- | --- |
| Ba1 | 3 | 3 | 0 | 3 | 1 | Piscivore |
| Ba2 | 6 | 4 | 1 | 4 | 3 | Piscivore |
| Ba3 | 4 | 3 | 0 | 3 | 1 | Piscivore |
| BuD | 3 | 3 | 0 | 4 | 2 | Planktivore |
| BuS | 3 | 3 | 0 | 1 | 2 | Planktivore |

## 4. Discussion

As coral reef communities change and reorganise in response to the increasing anthropogenic and climate disturbances, approaches that detect the new species configurations and their contribution to key ecosystem processes are required (Lester et al. 2020; Pombo-Ayora et al. 2020). Here, we show that the use of major categories (family level or above) in studying coral reef communities fails to identify strikingly distinct regimes. We also implement for the first time the use of a trait-based approach to study coral reef benthic communities, and show its relevance in the study and detection of different communities.

The spatio-temporal study of coral reefs is key to predicting their trajectories and recovery potential after disturbances (Ampou et al. 2017; Robinson et al. 2019). Within this context, global citizen science programs such as ReefCheck are vital for the temporal study of coral reefs, and contribute to community capacity building and education (Schäpply et al. 2017). Such programs, however, rely often on the study of coral communities using major categories (at family level for fishes and at class level or higher for benthic organisms), which as we show here might mask the presence of distinct assemblages. Given that current global change scenario is resulting in unprecedented ecosystem degradation, temporal monitoring of ecosystems and the detection of community changes is of foremost importance. However, the monitoring approaches might need to be readjusted or extended in order to provide higher taxonomic resolution surveys that capture the different emerging species configurations.

Changes in coral reefs communities and especially the decline of key reef building species contribute to the long-term functional erosion of coral reefs that could result in the loss of associated ecosystem services (Rogers et al. 2014; Hempson et al. 2017). For example, decreases in structural complexity and the associated loss of habitat structure have been associated with a decline in fish biomass and therefore fisheries (Rogers et al. 2014). However, as shown recently in some Caribbean reefs, not all communities with low-coral cover might display compromised ecosystem functioning (Lester et al. 2020), highlighting the need to understand the composition but also functioning of different coral reef communities. The use of trait-based approaches to gain insights into the role of biodiversity in ecosystem functioning has been successfully implemented to study and detect changes in fish and scleractinian coral communities (e.g. Darling et al. 2012; Mouillot et al. 2014), but to date such approaches have not yet been implemented to study coral reef benthic changes beyond scleractinian corals. Identification of coral reef benthic organisms to species or even genus level is extremely challenging, especially from visual census or imagery (Althaus et al. 2015), which probably has restrained researchers from applying trait-based approaches to whole coral reef benthic communities. Not only many groups of benthic organisms are highly understudied (Wee et al. 2017), but many other organisms such as sponges or soft corals require advanced genetic tools or microscopic examination for their taxonomic classification (McFadden et al. 2017; Ruiz et al. 2017; Koido et al. 2019). Here, we show that even if visual identification of many coral reef benthic organisms to species level remains impossible, the classification of organisms at the highest taxonomic level possible, which for some organisms may just be at the morphological level (i.e. sponges, Schönberg and Fromont, 2014) still yields high quality data on which trait-based approaches can be applied. Within our trait-based approach, in order to consider the inherent trait variability from categories that contain several species (e.g. massive sponges), we have used ordered categorical traits instead of continuous traits (e.g. growth ranges, broad lifespan ranges instead of specific values). The use of such an approach allowed us not only to delineate different community regimes that matched the ones identified using fish communities (at species level), but also to obtain insights into some of the functions that might be compromised in the different community regimes.

Our results show as previously highlighted by Ponti et al. (2016) that coral reef assemblages around Bangka and Bunaken islands are highly heterogeneous. More importantly, we observed that out of five different community regimes detected, only two were dominated by reef-building species, one of which was dominated by the blue coral *H. coerulea*. Dominance of the blue coral in other Indo-Pacific reefs has been previously reported and has been attributed to high growth, high thermal tolerance and its capacity of inhibiting scleractinian coral larval recruitment (Atrigenio et al. 2017; Guzman et al. 2019). Under the present scenario of climate change, communities dominated by *H. coerulea* might become increasingly common, but up to date, there is little information whereas *H. coerulea* dominated communities might sustain similar ecosystem functions as scleractinian dominated reefs (Guzman et al. 2019). Here, we show that the community dominated by *H. coerulea* (Ba2) presented comparable benthic and fish functional diversity to the scleractinian-dominated community (BuS). However, we would like to note that our surveys were one-time surveys and therefore a temporal data series is required to analyse the community trajectories and changes in functional diversity.

The approach used also allowed us to identify two communities that were dominated by potentially proliferating non-calcifying invertebrates displaying fast-growths and short lifespans. The proliferation of invertebrates able to overgrow live corals such as ascidians, sponges or some soft corals such as the opportunistic xeniids has been previously linked to the degradation of environmental conditions (Shenkar et al. 2008; Baum et al. 2016; Biggerstaff et al. 2017; Vollstedt et al. 2020). The dominance of benthic habitats by the soft coral *Xenia* spp. has been previously observed at different Indonesian reefs affected by blast fishing, including some reefs in Bunaken, Komodo and Wakatobi national parks (Fox et al. 2003; Marlow et al. 2019). Proliferation of colonial ascidians has also been reported in several reefs after increases in nutrient availability and overfishing (Shenkar et al. 2008; Roth et al. 2017; Tebett et al. 2019). Increases in these organisms can have severe effects on reef health and functioning by altering reef replenishment, geomorphology and trophic structure (Plass-Johnson et al. 2018; Tebett et al. 2019; Russ et al. 2020). Here, we observed that the two communities dominated by potentially proliferating organisms displayed simultaneously the lowest fish and benthic functional diversities, including the lowest diversity and cover of reef-building functional entities, such as branching corals. Many ascidians, soft corals and sponges possess varied chemical defences, that not only confer them spatial competitive advantages over corals, but that can also contribute to the inhibition of coral settlement, further contributing to coral loss (Atrigenio and Aliño 1996; Maida et al. 2001; Helber et al. 2018; Brandt et al. 2019). Proliferation of such organisms poses therefore a serious threat to the stability of coral reef ecosystems, however more studies are needed to further explore their implications on ecosystem functioning and their temporal persistence.

Sponges are the second most important invertebrate group (after corals) in determining substrate composition and nutrient cycles in coral reefs (de Goeij et al. 2013). Increasing evidence suggests that sponges might be increasing in abundance as consequence of climate change and anthropogenic stressors such as eutrophication, overfishing or sedimentation (Loh et al. 2015; Biggerstaff et al. 2018; Chaves-Fonnegra et al. 2018). Whereas the temporal persistence of sponge reefs and their functioning is still unclear and the subject of much current discussion (Lesser and Slattery, 2020; Pawlik and McMurray, 2020), an impeding limitation in hitherto existing studies is potentially the consideration of sponge reefs as a homogeneous entity. In fact, sponge species display marked divergence in their morphotypes and life histories, with some encrusting sponges such as *Terpios hoshinota* displaying turnover rates of months, whereas barrel sponges live thousands of years (Yomogida et al. 2017; McGrawth et al. 2019). Therefore, a sponge reef dominated by fast-growing short-lived sponges such as encrusting or bioeroding sponges might function completely different than a sponge reef composed mainly of slow-growing long-lived sponges, highlighting the need for the detailed study of such habitats and organisms functions. Here, all the sites surveyed at 10 m in the island of Bunaken were dominated by sponge communities. Interestingly, when the composition was studied using the taxonomic categories, we identified encrusting sponges as the major taxonomic benthic group in this cluster. However, when we explored the functional diversity and identified the most often encountered functional traits (CWM), we were able to observe that this community was in fact characterised by long-lived, slow-growing, massive, filter-feeder organisms, which included both massive and barrel sponges. These results demonstrate the importance of studying communities at the functional level, since they it enables a much more comprehensive understanding of the community and their ecosystem functioning (e.g. by considering similar traits from different taxonomic groups) than taxonomic categories. We also observed that the sponge-reefs observed in Bunaken displayed the highest fish functional diversity, the second highest benthic functional diversity, and contained most of the benthic functional entities contributing to reef building and accretion. Although, as mentioned earlier more surveys both in space and time are required to draw solid conclusions on the trajectories of coral reef communities around Bunaken and Bangka islands, our results suggest that the sponge-reefs identified might not be related to changing environmental conditions, but rather to other inherent reef characteristics such as topography (i.e. all the sponge-reef sites were reef walls).

In summary, by using a case study on coral reefs at the epicentre of tropical marine biodiversity we provide new evidence on 1) the importance of using high resolution taxonomic data for the detection of community regimes, 2) the usefulness of using trait-based approaches to explore and identify different community compositions and their contribution to key ecosystem processes and 3) the necessity of considering all benthic organisms to detect species configurations that are not dominated by scleractinian corals. Finally, we would like to highlight that although this study represents a successful example for the implementation of a trait-based approach on groups of similar benthic organisms, there is still a strong need for studies which provide trait-based knowledge on many of the understudied benthic organisms, in order to keep improving our knowledge of coral reef communities and improve the trait-based approaches used.

## Acknowledgement

We would like to thank Sam Ratulangi University (UNSRAT, Manado, Indonesia) staff from the Faculty of Fisheries and Marine Science, in particular Dr. Jane Mamuaja and everyone who supported the research and visa application. In addition, we would like to thank the staff from Bunaken Divers, Bunaken Island and the staff from Coral Eye, Bangka Island, who assisted us in carrying out the fieldwork. Research was carried out under LOA with Sam Ratulangi University, Manado, Indonesia from 14.1.2020, SIMAKSI number 246 from 18.2.2020 (to access and perform research around Bunaken Island) and with the research permits from the Ministry of Research, Technology, and Higher Education in Indonesia to PJS (n° 086/FRP/E5/Dit.KI/I/2020) and MR (n° 62/E5/E5.4/SIP/2020). Samples, photos and video material were transferred according to the Material Transfer Agreement between the University of Oldenburg (Germany) and the Sam Ratulangi University, Manado (Indonesia) from 18.11.2019. This research was funded by an Alexander von Humboldt post-doctoral fellowship to MR.

