## Supplementary material (Methods, Figures, Tables) for "Beyond coral-algal regimes: high taxonomic resolution surveys and trait-based analyses reveal multiple benthic regimes"

**Supplementary methods** **2**

Benthic transects and categories 2

Benthic trait categories 2

Fish trait categories 3

References 3

**Supplementary figures** **4**

Supplementary Figure 1 4

Supplementary Figure 2 5

**Supplementary Tables** **6**

Supplementary Table 1 6

Supplementary Table 2 7

Supplementary Table 3 16

#### Supplementary methods

*Benthic transects and categories*

The following major categories were used: hard coral, soft coral, sponges, crustose coralline algae, macroalgae, fleshy algae and other living organisms.

The high taxonomic resolution categories were chosen to ensure a reliable identification based uniquely on picture analysis. Scleractinian corals were mostly identified to genus level according to Veron et al. (2000), Budd et al. (2012) and Wallace et al. (2012). Some scleractinian coral genus such as *Porites*, *Montipora* and *Hynophora* that include species with marked morphologies and life histories (Darling et al. 2012) were further classified according to their morphologies (branching vs. massive/submassive forms). Soft corals were classified at genus level when possible. For example, the most abundant Alyconiidae genus *Klyxum/Cladiella*, *Lobophytum*, *Sarcophyton* and *Sinularia* were identified. However, xenid corals that often require detailed morphological analysis to determine their genus were studied at family level (Fabricious and Alderslade 2001). Most Neptheidae were identified at family level, with only the conspicuous *Dendronephthya* being identified at genus level. Corallimorpharians and sea anemones were identified at order level (Actinaria, Corallimorpharia). The most common zoanthid (*Palythoa* spp.) was identified, whereas the rest were classified at order level. The most common and conspicuous hydroids encountered were identified at genus or species level (e.g *Aglaophenia cupressina*, *Myrionema* spp.), whereas the rest were identified at class level. The hydrocoral *Millepora* was identified as genus level, but it was included into the hard coral major category, instead of other living animals as the rest of hydroids. Sponge categories, were based on morphological classification, as this has been previously suggested as an appropriate proxy of sponge species diversity (Bell and Barnes 2001; Berman et al. 2013). Only the encrusting species *Lamellodysidea herbacea* was identified at species level, since this species is conspicuous and can be the dominant benthic organisms in Indonesian reefs (Bigerstaff et al. 2017). Ascidians were initially separated into solitary and colonial species, with some particularly relevant (e.g. abundant) and conspicuous species or families being identified (e.g. Didemnidae, *Polycarpa aurata*). Crustose coralline algae were separated into two morphs (encrusting and articulated). The most common genus of macroalgae were also identified: *Turbinaria*, *Tydemania* and *Halimeda*.

*Benthic trait categories*

The functional ecology of benthos categories was characterised using 12 traits: colony formation, growth form, maximum colony size, longevity, growth rate, body flexibility, skeleton presence, reproductive strategy, sexual system, feeding strategy, presence of photosynthetic symbionts and corallite maximum width (only for scleractinian corals). Colony formation was coded as binary ("0" solitary organisms, "1" colonial/multimodal organism). Growth form was classified into three categories: "massive", "encrusting" or "branching". Maximum colony size was coded using three ordered categories: "1" for colonies/individuals smaller than 10 cm, "2" for organisms between 10.1-100 cm and "3" for organisms that can grow beyond 100 cm. In case of organisms growing asymmetrically, the direction of maximum growth (e.g. vertical) was considered. Longevity was coded using four ordered categories: "1" for organisms living < 1 year, "2" for organisms living between 1-10 years, "3" for organisms living between 10-50 years and "4" for organisms with a lifespan > 50 years. Growth rate was coded using three ordered categories: "1" for organisms growing up to 3 cm/year (in the direction of their maximum growth), "2" for organisms with growths of 3.1-10 cm/year and "3" with organisms growing more than 10 cm/year. Body flexibility was coded using three ordered categories: "not flexible", "limited flexibility (≤ 45°)", "highly flexible (> 45°)". Skeleton presence was coded as binary ("0" for organisms without a hard exoskeleton and "1" for organisms with a skeleton). Reproductive strategy was coded as binary ("0" for brooders and "1" for broadcasters).

Adult sexual distribution was classified in three categories "hermaphrodites", "gonochoric" and "alternate" which referred to organism categories containing adults with both distributions (e.g. some sponges or algae). Feeding strategy was classified into three categories: "photosynthetic", "filter-feeder" and "selective" which was attributed to organisms that selectively captured their prey with tentacles, for example. Since the presence of photosynthetic symbionts is an important attribute that is often species-specific we used three ordered categories that consider species variability within a benthic category: "1" for species groups that never host photosynthetic symbionts, "2" for benthic groups where photosynthetic symbionts sometimes present and "3" for species/groups that always harbour photosynthetic symbionts. Corallite maximum width was classified into five ordered categories: "1" for organisms lacking corallites, "2" for corallites smaller than 2 mm, "3" for corallites between 2-10 mm, "4" for corallites between 10.1-20 mm and "5" for corallites bigger than 20 mm.

*Fish trait categories*

The functional ecology of fish species was characterised using six traits (body size, diet, period of activity, vertical position, gregariousness and mobility). Fish body size was coded using six ordered categories: 0-7 cm, 7.1-15 cm, 15.1-30 cm, 30.1-50 cm, 50.1-80 cm and > 80 cm. Diet was characterised based on main items consumed by each species following earlier classifications (MacNeil et al. 2015; Bierwagen et al. 2018), which led to eight trophic categories: "browsers" (i.e. fish eating macroalgae), "grazers/detritivores" (i.e. fish feeding on turf algae or particulate organic material), "excavators/scrapers" (i.e. fish that feed on algae and scrape or excavate the reef substratum), "corallivores" (i.e. fish that feed strictly on corals), "micro-invertivores" (i.e. fish that feed on small invertebrates), "macro-invertivores" (i.e. fish that feed on larger invertebrates), "omnivores" (i.e. fish that feed on a wide variety of material including plankton, algae and invertebrates), "planktivores" (i.e. fish eating small organisms in the water column) and "piscivores" (i.e. fish eating fish and large invertebrates such as cephalopods). The period of activity (i.e. the period at which fish feed) was coded as binary with "0" for diurnal species and "1" for nocturnal species. The vertical position of fish in the water column was coded using three ordered categories: "1": benthic, "2": bentho-pelagic and "3" "pelagic". Gregariousness (i.e. schooling behaviour) was coded using four ordered categories: "1" solitary, "2" pairing, "3" living in small groups (< 20 individuals) and "4" schooling species (> 20 individuals). Mobility was coded using four ordered categories following Olivier et al. (2018): "1" highly site-attached species (i.e. territorial species)", "2" mobile species with a small home range, "3" mobile species with a large home range and "4" widely mobile species with a very large home range (e.g. species that can travel very long distances such as Carangidae).

*References*


Bell, J., and D. Barnes. 2001. Sponge morphological diversity: A qualitative predictor of species diversity? Aquatic Conservation: Marine and Freshwater Ecosystems 11:109-121.

Berman, J., M. Burton, R. Gibbs, K. Lock, P. Newman, J. Jones, and J. Bell. 2013. Testing the suitability of a morphological monitoring approach for identifying temporal variability in a temperate sponge assemblage. Journal for Nature Conservation 21:173-182.

Biggerstaff, A., J. Jompa, and J. Bell. 2017. Increasing benthic dominance of the phototrophic sponge *Lamellodysidea herbacea* on a sedimented reef within the Coral Triangle. Marine Biology 164.

Budd AF, Fukami H, Smith ND, Knowlton N. 2012. Taxonomic classification of the reef coral family Mussidae (Cnidaria: Anthozoa: Scleractinia). Zoological Journal of the Linnean Society 166(3):465-529.

Darling, E. S., L. Alvarez-Filip, T. A. Oliver, T. R. McClanahan, and I. M. Côté. 2012. Evaluating life-history strategies of reef corals from species traits. Ecology Letters 15:1378-1386.

Veron, J. E. N. 2000. Corals of the World. Townsville: Australian Institute of Marine Science.

Wallace CC, Done BJ, Muir PR. 2012. Revision and catalogue of worldwide staghorn corals *Acropora* and *Isopora* (Scleractina: Acroporidae) in the Museum of Tropical Queensland. South Brisbane: Queensland Museum.

### Supplementary figures

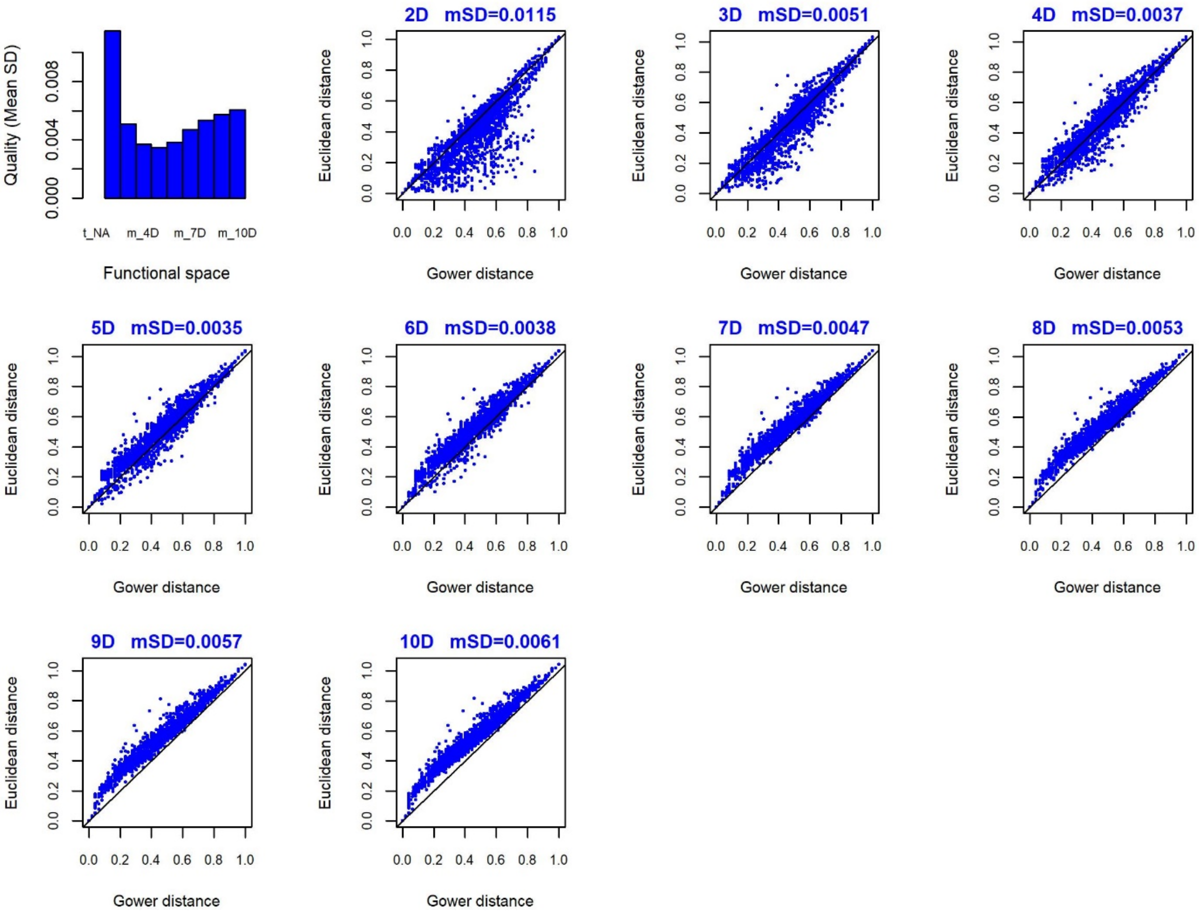


**Supplementary figure 1.** Quality of the different benthic functional spaces created (up to 10 axis) and their mean squared deviations (mSD).

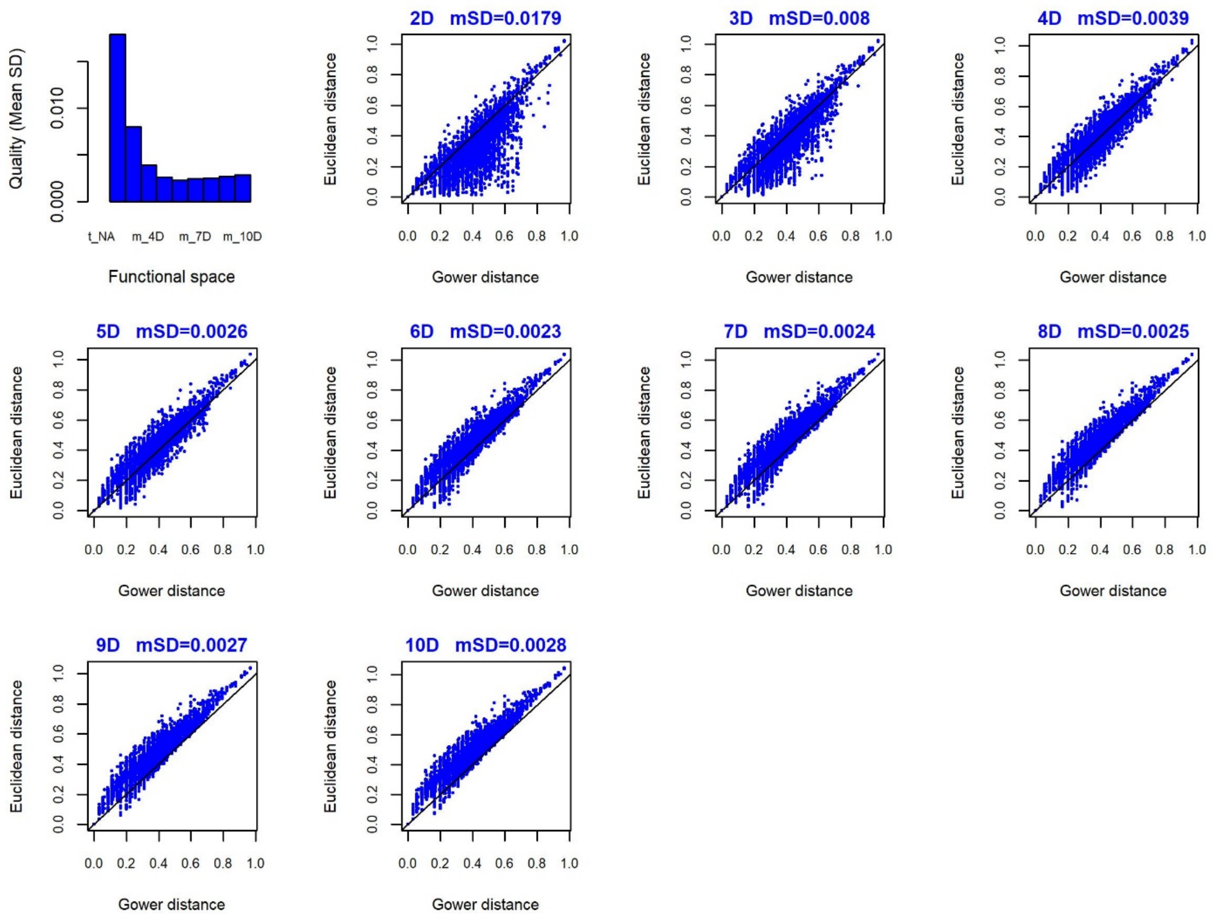


**Supplementary figure 2**. Quality of the different fish functional spaces created (up to 10 axis) and their mean squared deviations (mSD).

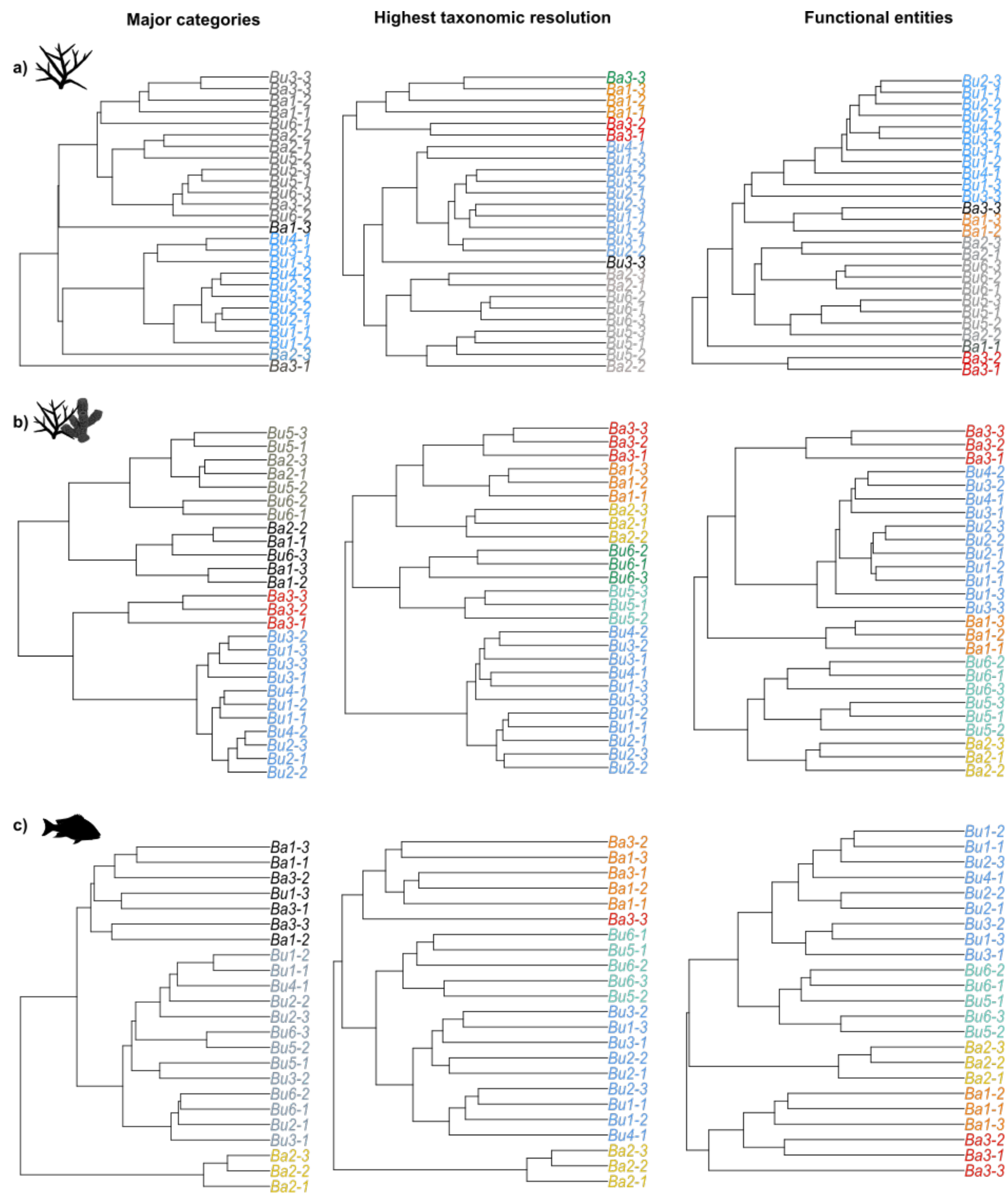


**Supplementary figure 2**. Cluster analysis showing the site similarities by studying the coral communities (a), benthic communities (corals and other organisms, b) and fish communities (c) of using major categories, high taxonomic resolution categories and functional entities (FEs).

#### Supplementary Tables

**Supplementary table 1:** Main characteristics of the study sites.

| Island | Code | Site | Latitude | Longitude | Island site | Depth | Habitat | Marine Protected Area |
|---|---|---|---|---|---|---|---|---|
| Bunaken | Bu1 | Mandolin | 1.611474° | 124.7328° | W | 10 | Wall | Y |
| | Bu2 | Tengah | 1.615678° | 124.7322° | W | 10 | Wall | Y |
| | Bu3 | Likupang II | 1.60002° | 124.7668° | SW | 10 | Wall | Y |
| | Bu4 | Likupang III | 1.603537° | 124.7662° | SW | 10 | Wall | Y |
| | Bu5 | Timur 3 | 1.612168° | 124.7829° | E | 3 | Outer lagoon | Y |
| | Bu6 | Timur 2 | 1.607365° | 124.7823° | E | 3 | Outer lagoon | Y |
| Bangka | Ba1 | Coral eye | 1.750501° | 125.1334° | SW | 10 | Gentle slope | N |
| | Ba3 | Sipi | 1.78582° | 125.1303° | W | 10 | Gentle slope | N |
| | Ba2 | Coral Eye permanent | 1.748881° | 125.1337° | SW | 3 | Lagoon | N |

**Supplementary table 2**. Details on the different benthic categories identified (at the highest taxonomic resolution possible: Unique_id). Categories having the same FE (Functional Entity number) grouped as one functional entity. I/C: colonial or individual organisms (0: individual, 1: colonial or modular). BF: body form (B: branching, M: massive, E: encrusting). F: flexibility (1: no flexibility, 2: limited flexibility <45°, 3: high flexibility >45°). FS: feeding strategy (FF: filter feeder, Phot: photosynthesis, Sel: selective feeder). PS: presence of photosynthetic symbionts (1: none, 2: sometimes, 3: always). RS: reproductive strategy (Alt: alternate, Brood: brooder, Broad: broadcaster). AR: adult reproductive organs (H: hermaphrodite, G: gonochoric, O: other). S: exoskeleton (0: no, 1: yes). CMW: corallite maximum width (1: no corallites, 2: < 2 mm, 3: 2-10 mm, 4: 10.1-20 mm, 5: > 20 mm). GR: growth rate (1: < 3 cm/year, 2: 3.1-10 cm/year, 3: > 10 cm/year). CS: colony maximum size (1: <10 cm, 2: 10.1-100 cm, 3: > 100 cm), L: longevity (1: < 1 year, 2: 1-10 years, 3:10-50 years, 4: > 50 years). Ref: references.

| Unique id | Major category | FE | I/C | BF | F | FS | PS | RS | AR | S | CMW | GR | CS | L | Ref. |
|---|---|---|---|---|---|---|---|---|---|---|---|---|---|---|---|
| **CCA _articulate** | Coralline algae | #1 | 0 | B | 2 | Phot. | 1 | Alt. | O | 1 | 1 | 2 | 1 | 3 | 30-32 |
| ***Polycarpa_aurata*** | Other live | #10 | 0 | M | 2 | FF | 1 | Brood. | H | 0 | 1 | 3 | 1 | 1 | 19 |
| **Ascidian_solitary** | Other live | #11 | 0 | M | 2 | FF | 2 | Brood. | H | 0 | 1 | 3 | 1 | 1 | 19 |
| **Actinaria** | Other live | #12 | 0 | M | 3 | Sel. | 2 | Broadc. | G | 0 | 1 | 3 | 2 | 2 | 2-5 |
| ***Millepora*_spp** | Hard coral | #13 | 1 | B | 1 | Sel. | 3 | Broadc. | G | 1 | 1 | 2 | 3 | 4 | 91 |
| ***Porites*_branching_spp** | Hard coral | #14 | 1 | B | 1 | Sel. | 3 | Broadc. | G | 1 | 2 | 2 | 3 | 4 | 1 |
| ***Pavona*_spp** | Hard coral | #15 | 1 | B | 1 | Sel. | 3 | Broadc. | G | 1 | 3 | 1 | 3 | 4 | 1 |
| ***Acropora*_spp.** | Hard coral | #16 | 1 | B | 1 | Sel. | 3 | Broadc. | H | 1 | 2 | 2 | 3 | 4 | 1 |
| ***Montipora*_branching_sp** | Hard coral | #16 | 1 | B | 1 | Sel. | 3 | Broadc. | H | 1 | 2 | 2 | 3 | 4 | 1 |
| ***Pocillopora_damicornis*** | Hard coral | #16 | 1 | B | 1 | Sel. | 3 | Broadc. | H | 1 | 2 | 2 | 3 | 4 | 1 |
| ***Pocillopora*_spp** | Hard coral | #16 | 1 | B | 1 | Sel. | 3 | Broadc. | H | 1 | 2 | 2 | 3 | 4 | 1 |
| **Hydnophora_branching_spp** | Hard coral | #17 | 1 | B | 1 | Sel. | 3 | Broadc. | H | 1 | 3 | 2 | 3 | 4 | 1 |
| **Heliopora_coerulea** | Hard coral | #18 | 1 | B | 1 | Sel. | 3 | Brood. | G | 1 | 1 | 2 | 3 | 4 | 81-83 |
| ***Seriatopora*_spp** | Hard coral | #19 | 1 | B | 1 | Sel. | 3 | Brood. | H | 1 | 2 | 1 | 2 | 4 | 1 |
| ***Stylophora*_spp** | Hard coral | #19 | 1 | B | 1 | Sel. | 3 | Brood. | H | 1 | 2 | 1 | 2 | 4 | 1 |
| ***Turbinaria*_algae_spp** | Macroalgae | #2 | 0 | B | 3 | Phot. | 1 | Alt. | O | 0 | 1 | 3 | 1 | 1 | 116, 117 |
| ***Isopora*_spp** | Hard coral | #20 | 1 | B | 1 | Sel. | 3 | Brood. | H | 1 | 2 | 1 | 3 | 4 | 1 |
| **Alcyoniidae_other_spp** | Soft coral | #21 | 1 | B | 2 | Sel. | 2 | Broadc. | G | 0 | 1 | 3 | 3 | 3 | 9-13 |
| ***Lobophytum*_spp** | Soft coral | #21 | 1 | B | 2 | Sel. | 2 | Broadc. | G | 0 | 1 | 3 | 3 | 3 | 13, 85-86 |
| **Arborescent_sponge** | Sponge | #22 | 1 | B | 3 | FF | 2 | Alt. | H | 0 | 1 | 2 | 2 | 3 | 14-18 |
| **Repent_sponge** | Sponge | #22 | 1 | B | 3 | FF | 2 | Alt. | H | 0 | 1 | 2 | 2 | 3 | 14-18, 102 |
| **Erect_sponge** | Sponge | #22 | 1 | B | 3 | FF | 2 | Broadc. | H | 0 | 1 | 2 | 2 | 3 | 14-18 |
| ***Dendronephtya*_spp** | Soft coral | #23 | 1 | B | 3 | Sel. | 1 | Brood. | G | 0 | 1 | 3 | 2 | 3 | 9, 10, 13, 43, 44 |
| **Hydroid_others** | Other live | #24 | 1 | B | 3 | Sel. | 2 | Broadc. | G | 0 | 1 | 3 | 1 | 1 | 6-8 |
| **Soft_coral_other_spp** | Soft coral | #25 | 1 | B | 3 | Sel. | 2 | Broadc. | G | 0 | 1 | 3 | 3 | 3 | 9, 13, 111-115 |

| | | | | | | | | | | | | | | | |
|---|---|---|---|---|---|---|---|---|---|---|---|---|---|---|---|
| **Sea_fans** | Soft coral | #26 | 1 | B | 3 | Sel. | 2 | Brood. | G | 0 | 1 | 2 | 3 | 3 | 13, 72-77 |
| **Nephteiidae_other_spp** | Soft coral | #27 | 1 | B | 3 | Sel. | 2 | Brood. | G | 0 | 1 | 3 | 2 | 3 | 9, 10, 13, 92-94 |
| ***Aglaophenia_cupressina*** | Other live | #28 | 1 | B | 3 | Sel. | 3 | Broadc. | G | 0 | 1 | 3 | 2 | 1 | 6-8 |
| ***Klyxum***_**spp** | Soft coral | #29 | 1 | B | 3 | Sel. | 3 | Broadc. | G | 0 | 1 | 3 | 2 | 3 | 9-13 |
| **Macroalgae_other_spp** | Macroalgae | #3 | 0 | B | 3 | Phot. | 1 | Alt. | O | 0 | 1 | 3 | 2 | 2 | 85-88 |
| ***Tydemania***_**spp** | Macroalgae | #3 | 0 | B | 3 | Phot. | 1 | Alt. | O | 0 | 1 | 3 | 2 | 2 | 117 |
| ***Sinularia***_**spp** | Hard coral | #30 | 1 | B | 3 | Sel. | 3 | Broadc. | G | 0 | 1 | 3 | 3 | 4 | 9, 13, 104, 107-110 |
| **Bryozoan** | Other live | #31 | 1 | E | 1 | FF | 1 | Alt. | H | 1 | 1 | 2 | 2 | 2 | 22-26 |
| ***Celleporaria_sibogae*** | Other live | #31 | 1 | E | 1 | FF | 1 | Brood. | H | 1 | 1 | 2 | 2 | 2 | 22-26, 34 |
| **Encrusting_sponge** | Sponge | #32 | 1 | E | 1 | FF | 2 | Alt. | H | 0 | 1 | 3 | 3 | 1 | 53-58 |
| **Ascidian_colonial** | Other live | #33 | 1 | E | 1 | FF | 2 | Brood. | H | 0 | 1 | 3 | 2 | 2 | 19 |
| **Encrusting_ascidian** | Other live | #33 | 1 | E | 1 | FF | 2 | Brood. | H | 0 | 1 | 3 | 2 | 2 | 19, 50-52 |
| **Didemnidae_incrusting** | Other live | #34 | 1 | E | 1 | FF | 3 | Brood. | H | 0 | 1 | 3 | 2 | 2 | 19, 46-49 |
| ***Lissoclinum***_**spp** | Other live | #34 | 1 | E | 1 | FF | 3 | Brood. | H | 0 | 1 | 3 | 2 | 2 | 19, 46-49 |
| ***Lamelodysea_herbacea*** | Sponge | #35 | 1 | E | 1 | FF | 3 | Brood. | H | 0 | 1 | 3 | 3 | 1 | 84 |
| **Zoanthid_other_spp** | Other live | #36 | 1 | E | 1 | Sel. | 2 | Broadc. | H | 0 | 1 | 2 | 2 | 3 | 96-100 |
| **Hard_coral_massive_encrusting_other** | Hard coral | #37 | 1 | E | 1 | Sel. | 3 | Broadc. | G | 1 | 2 | 1 | 3 | 4 | 1 |
| ***Coscinaraea***_**spp** | Hard coral | #38 | 1 | E | 1 | Sel. | 3 | Broadc. | G | 1 | 3 | 1 | 3 | 4 | 1 |
| ***Leptastrea***_**spp** | Hard coral | #38 | 1 | E | 1 | Sel. | 3 | Broadc. | G | 1 | 3 | 1 | 3 | 4 | 1 |
| ***Leptoseris***_**spp** | Hard coral | #38 | 1 | E | 1 | Sel. | 3 | Broadc. | G | 1 | 3 | 1 | 3 | 4 | 1 |
| ***Pachyseris***_**spp** | Hard coral | #38 | 1 | E | 1 | Sel. | 3 | Broadc. | G | 1 | 3 | 1 | 3 | 4 | 1 |
| ***Turbinaria***_**coral_spp** | Hard coral | #38 | 1 | E | 1 | Sel. | 3 | Broadc. | G | 1 | 3 | 1 | 3 | 4 | 1 |
| ***Palythoa***_**spp** | Other live | #39 | 1 | E | 1 | Sel. | 3 | Broadc. | H | 0 | 1 | 2 | 2 | 3 | 96-100 |
| ***Halimeda***_**spp** | Macroalgae | #4 | 0 | B | 3 | Phot. | 1 | Broadc. | O | 0 | 1 | 3 | 1 | 2 | 78-80 |
| ***Echinopora***_**spp** | Hard coral | #40 | 1 | E | 1 | Sel. | 3 | Broadc. | H | 1 | 3 | 1 | 3 | 4 | 1 |
| ***Merulina***_**spp** | Hard coral | #40 | 1 | E | 1 | Sel. | 3 | Broadc. | H | 1 | 3 | 1 | 3 | 4 | 1 |
| ***Mycedium***_**spp** | Hard coral | #40 | 1 | E | 1 | Sel. | 3 | Broadc. | H | 1 | 3 | 1 | 3 | 4 | 1 |
| ***Oxypora***_**spp** | Hard coral | #40 | 1 | E | 1 | Sel. | 3 | Broadc. | H | 1 | 3 | 1 | 3 | 4 | 1, 95 |
| ***Echinophyllia***_**spp** | Hard coral | #41 | 1 | E | 1 | Sel. | 3 | Broadc. | H | 1 | 4 | 1 | 3 | 4 | 1 |
| ***Pectinia***_**spp** | Hard coral | #41 | 1 | E | 1 | Sel. | 3 | Broadc. | H | 1 | 4 | 1 | 3 | 4 | 1, 101 |
| ***Moseleya***_**spp** | Hard coral | #42 | 1 | E | 1 | Sel. | 3 | Broadc. | H | 1 | 5 | 1 | 2 | 4 | 1 |
| ***Myrionema***_**spp** | Other live | #43 | 1 | E | 2 | Sel. | 3 | Brood. | G | 0 | 1 | 3 | 2 | 1 | 6-8 |
| **Fleshy_sponge** | Sponge | #44 | 1 | M | 1 | FF | 2 | Alt. | H | 0 | 1 | 2 | 2 | 3 | 18, 63 |
| **Massive_sponge** | Sponge | #45 | 1 | M | 1 | FF | 2 | Alt. | H | 0 | 1 | 2 | 3 | 3 | 61, 69, |

| | | | | | | | | | | | | | | | |
|---|---|---|---|---|---|---|---|---|---|---|---|---|---|---|---|
| | | | | | | | | | | | | | | | 89-90 |
| **Barrel_sponge** | Sponge | #46 | 1 | M | 1 | FF | 2 | Broadc. | G | 0 | 1 | 1 | 3 | 4 | 18, 20, 21 |
| **Globular_sponge** | Sponge | #47 | 1 | M | 1 | FF | 2 | Broadc. | H | 0 | 1 | 2 | 2 | 3 | 69-71 |
| **Calcareous_sponge** | Sponge | #48 | 1 | M | 1 | FF | 2 | Brood. | H | 0 | 1 | 2 | 2 | 3 | 27-29 |
| ***Tubastrea*****_spp** | Hard coral | #49 | 1 | M | 1 | Sel. | 1 | Brood. | G | 1 | 1 | 1 | 3 | 4 | 1 |
| **CCA_encrusting** | Coralline algae | #5 | 0 | E | 1 | Phot. | 1 | Alt. | O | 1 | 1 | 1 | 2 | 4 | 30-33 |
| ***Porites*****_other_spp** | Hard coral | #50 | 1 | M | 1 | Sel. | 3 | Broadc. | G | 1 | 2 | 1 | 3 | 4 | 1 |
| ***Diploastrea*****_spp** | Hard coral | #51 | 1 | M | 1 | Sel. | 3 | Broadc. | G | 1 | 3 | 1 | 3 | 4 | 1 |
| ***Galaxea*****_spp** | Hard coral | #51 | 1 | M | 1 | Sel. | 3 | Broadc. | G | 1 | 3 | 1 | 3 | 4 | 1 |
| ***Gardinoserosis*****_spp** | Hard coral | #51 | 1 | M | 1 | Sel. | 3 | Broadc. | G | 1 | 3 | 1 | 3 | 4 | 1 |
| ***Goniopora*****_spp** | Hard coral | #51 | 1 | M | 1 | Sel. | 3 | Broadc. | G | 1 | 3 | 1 | 3 | 4 | 1 |
| ***Euphyllia*****_spp** | Hard coral | #52 | 1 | M | 1 | Sel. | 3 | Broadc. | G | 1 | 5 | 1 | 3 | 4 | 1 |
| ***Plerogyra*****_spp** | Hard coral | #52 | 1 | M | 1 | Sel. | 3 | Broadc. | G | 1 | 5 | 1 | 3 | 4 | 1 |
| ***Montipora*****_submassive** | Hard coral | #53 | 1 | M | 1 | Sel. | 3 | Broadc. | H | 1 | 2 | 1 | 3 | 4 | 1 |
| ***Favia*****_spp** | Hard coral | #54 | 1 | M | 1 | Sel. | 3 | Broadc. | H | 1 | 3 | 1 | 2 | 4 | 1 |
| ***Alveopora*****_spp** | Hard coral | #55 | 1 | M | 1 | Sel. | 3 | Broadc. | H | 1 | 3 | 1 | 3 | 4 | 1 |
| ***Astreopora*****_spp** | Hard coral | #55 | 1 | M | 1 | Sel. | 3 | Broadc. | H | 1 | 3 | 1 | 3 | 4 | 1 |
| ***Cyphastrea*****_spp** | Hard coral | #55 | 1 | M | 1 | Sel. | 3 | Broadc. | H | 1 | 3 | 1 | 3 | 4 | 1 |
| ***Goniastrea*****_spp** | Hard coral | #55 | 1 | M | 1 | Sel. | 3 | Broadc. | H | 1 | 3 | 1 | 3 | 4 | 1 |
| ***Hydnophora*****_other_spp** | Hard coral | #55 | 1 | M | 1 | Sel. | 3 | Broadc. | H | 1 | 3 | 1 | 3 | 4 | 1 |
| ***Leptoria*****_spp** | Hard coral | #55 | 1 | M | 1 | Sel. | 3 | Broadc. | H | 1 | 3 | 1 | 3 | 4 | 1 |
| ***Phymastrea*****_spp** | Hard coral | #55 | 1 | M | 1 | Sel. | 3 | Broadc. | H | 1 | 3 | 1 | 3 | 4 | 1 |
| ***Platygyra*****_spp** | Hard coral | #55 | 1 | M | 1 | Sel. | 3 | Broadc. | H | 1 | 3 | 1 | 3 | 4 | 1 |
| ***Acanthastrea*****_spp.** | Hard coral | #56 | 1 | M | 1 | Sel. | 3 | Broadc. | H | 1 | 4 | 1 | 3 | 4 | 1 |
| ***Caulastrea*****_spp** | Hard coral | #56 | 1 | M | 1 | Sel. | 3 | Broadc. | H | 1 | 4 | 1 | 3 | 4 | 1 |
| ***Favites*****_spp** | Hard coral | #56 | 1 | M | 1 | Sel. | 3 | Broadc. | H | 1 | 4 | 1 | 3 | 4 | 1 |
| ***Oulophyllia*****_spp** | Hard coral | #56 | 1 | M | 1 | Sel. | 3 | Broadc. | H | 1 | 4 | 1 | 3 | 4 | 1 |
| ***Symphillia*****_spp** | Hard coral | #57 | 1 | M | 1 | Sel. | 3 | Broadc. | H | 1 | 5 | 1 | 2 | 4 | 1 |
| ***Lobophyllia*****_spp** | Hard coral | #58 | 1 | M | 1 | Sel. | 3 | Broadc. | H | 1 | 5 | 1 | 3 | 4 | 1 |
| ***Nephtheis_fascicularis*** | Other live | #59 | 1 | M | 2 | FF | 1 | Brood. | H | 0 | 1 | 3 | 1 | 2 | 19 |
| **Foraminifera** | Other live | #6 | 0 | E | 1 | Sel. | 2 | Alt. | G | 1 | 1 | 2 | 1 | 2 | 64-66 |
| **Papillate_sponge** | Sponge | #60 | 1 | M | 2 | FF | 2 | Broadc. | H | 0 | 1 | 2 | 2 | 3 | 61, 69, 89-90 |
| **Pedunculate_sponge** | Sponge | #60 | 1 | M | 2 | FF | 2 | Broadc. | H | 0 | 1 | 2 | 2 | 3 | 69-71 |
| ***Didemnum_molle*** | Other live | #61 | 1 | M | 2 | FF | 3 | Brood. | H | 0 | 1 | 3 | 2 | 2 | 19, 40-42 |
| ***Sarcophyton*****_spp** | Soft coral | #62 | 1 | M | 2 | Sel. | 3 | Broadc. | G | 0 | 1 | 2 | 3 | 3 | 9, 13, 103-106 |
| **Flabellate_sponge** | Sponge | #63 | 1 | M | 3 | FF | 2 | Alt. | G | 0 | 1 | 3 | 2 | 3 | 59-62 |
| **Xeniidae** | Soft coral | #64 | 1 | M | 3 | Sel. | 3 | Brood. | H | 0 | 1 | 3 | 1 | 2 | 9, 13, 119-122 |

| | | | | | | | | | | | | | | | |
|---|---|---|---|---|---|---|---|---|---|---|---|---|---|---|---|
| **Corallimorpharian** | Other live | #7 | 0 | E | 1 | Sel. | 2 | Broadc. | G | 0 | 1 | 3 | 2 | 2 | 35-38 |
| **Cyanobacteria** | Fleshy algae | #8 | 0 | E | 2 | Phot. | 1 | Alt. | O | 0 | 1 | 3 | 2 | 1 | 39 |
| **Turf** | Fleshy algae | #8 | 0 | E | 2 | Phot. | 1 | Alt. | O | 0 | 1 | 3 | 2 | 1 | 118 |
| ***Fungia*_spp** | Hard coral | #9 | 0 | M | 1 | Sel. | 3 | Broadc. | G | 1 | 5 | 1 | 2 | 3 | 1, 67-68 |

*References*


1. Madin, J. S., K. D. Anderson, M. H. Andreasen, T. C. L. Bridge, S. D. Cairns, S. R. Connolly, E. S. Darling, M. Diaz, D. S. Falster, E. C. Franklin, R. D. Gates, A. M. T. Harmer, M. O. Hoogenboom, D. Huang, S. A. Keith, M. A. Kosnik, C.-Y. Kuo, J. M. Lough, C. E. Lovelock, O. Luiz, J. Martinelli, T. Mizerek, J. M. Pandolfi, X. Pochon, M. S. Pratchett, H. M. Putnam, T. E. Roberts, M. Stat, C. C. Wallace, E. Widman, and A. H. Baird. 2016. The Coral Trait Database, a curated database of trait information for coral species from the global oceans. Scientific Data 3:160017.
2. Dixon, A. K., M. J. McVay, and N. E. Chadwick. 2017. Demographic modelling of giant sea anemones: population stability and effects of mutualistic anemonefish in the Jordanian Red Sea. Marine and Freshwater Research 68:2145-2155.
3. McVay, M. J. 2015. Population dynamics of clownfish sea anemones: Patterns of decline, symbiosis with anemonefish, and management for sustainability.
4. O'Reilly, E., B. M. Titus, M. W. Nelsen, S. Ratchford, and N. E. Chadwick. 2018. Giant ephemeral anemones? Rapid growth and high mortality of corkscrew sea anemones *Bartholomea annulata* (Le Sueur, 1817) under variable conditions. Journal of Experimental Marine Biology and Ecology 509:44-53.
5. Rifa i, M. A., Fatmawati, F. Tony, and H. Kudsiah. 2016. The survival and growth rate of three species of sea anemones from asexual reproduction in Pulau Kerumputan and Pulau Karayaan, Indonesia. Ecology, environment & conservation 22:1523-1531.
6. Boero, F. 1984. The ecology of marine hydroids and effects of environmental factors: A review. Marine Ecology 5:93-118.
7. Bosch-Belmar, M., A. Escurriola, G. Milisenda, V. L. Fuentes, and S. Piraino. 2019. Harmful fouling communities on fish farms in the SW Mediterranean sea: composition, growth and reproductive periods. Journal of Marine Science and Engineering 7:288.
8. Di Camillo, C.G., Bavestrello, G., Cerrano, C., Gravili, C., Piraino, S, Puce, S., Boero, F. 2017. Hydroids (Cnidaria, Hydrozoa): A neglected component of animal forests. Marine Animal Forests, 20, 397-427.
9. Fautin, D., J. Westfall, P. Cartwrigh, M. Daly, and C. Wyttenbach. 2004. Coelenterate Biology 2003: Trends in Research on Cnidaria and Ctenophora.
10. Benayahu, Y., and Y. Loya. 1987. Long-term recruitment of soft-corals (Octocorallia: Alcyonacea) on artificial substrata at Eilat (Red Sea). Marine Ecology Progress Series 38:161-167.
11. Goh, B. P. L., G. E. Tan, and L. T. Tan. 2011. Diversity, distribution and biological activity of soft corals (Octocorallia, Alcyonacea) in Singapore. Journal of Coastal Development 12:10.
12. Fitriani, D., M. Zainuri, and W. Nugraha. 2020. Laju Pertumbuhan dan Pertumbuhan Mutlak Karang Lunak Cladiella sp. pada Substrat yang Berbeda. Jurnal Kelautan Tropis 23:29.
13. Fabricius, K., Alderslade, P. 2001. Soft corals and sea fans: a comprehensive guide to the tropical shallow water genera of the central-west Pacific, the Indian Ocean and the Red Sea.: AIMS, Townsville, Qld. (Australia). 264 pp.
14. Lesser, M. P. 2006. Benthic–pelagic coupling on coral reefs: Feeding and growth of Caribbean sponges. Journal of Experimental Marine Biology and Ecology 328:277-288.
15. Schiefenhövel, K., and A. Kunzmann. 2012. Sponge farming trials: survival, attachment, and growth of two Indo-Pacific sponges, *Neopetrosia* sp. and *Stylissa massa*. Journal of Marine Biology 2012:417360.
16. Trautman, D. A., R. Hinde, and M. A. Borowitzka. 2000. Population dynamics of an association between a coral reef sponge and a red macroalga. Journal of Experimental Marine Biology and Ecology 244:87-105.

17. Mercado-Molina, A. E., A. M. Sabat, and P. M. Yoshioka. 2011. Demography of the demosponge *Amphimedon compressa*: Evaluation of the importance of sexual versus asexual recruitment to its population dynamics. Journal of Experimental Marine Biology and Ecology 407:355-362.
18. Erwin, P. M., and R. W. Thacker. 2007. Incidence and identity of photosynthetic symbionts in Caribbean coral reef sponge assemblages. Journal of the Marine Biological Association of the United Kingdom 87:1683-1692.
19. Millar, R. H. 1971. The Biology of Ascidians. Pages 1-100 *in* F. S. Russell and M. Yonge, editors. Advances in Marine Biology. Academic Press.
20. McMurray, S. E., J. E. Blum, and J. R. Pawlik. 2008. Redwood of the reef: growth and age of the giant barrel sponge *Xestospongia muta* in the Florida Keys. Marine Biology 155:159-171.
21. Fromont, J., and P. R. Bergquist. 1994. Reproductive biology of three sponge species of the genus *Xestospongia* (Porifera: Demospongiae: Petrosida) from the Great Barrier Reef. Coral Reefs 13:119-126.
22. Winston, J. E. 2017. Feeding behavior of modern bryozoans. Notes for a short course: Studies in Geology 5:1-21.
23. Winston, J. E., and J. B. C. Jackson. 1984. Ecology of cryptic coral reef communities. IV. Community development and life histories of encrusting cheilostome bryozoa. Journal of Experimental Marine Biology and Ecology 76:1-21.
24. Smith, A. M. 2014. Growth and calcification of marine bryozoans in a changing ocean. The Biological Bulletin 226:203-210.
25. Nielsen, C. and Larwood, G. P. (1985) Bryozoa : Ordovician to recent : papers presented at the 6th International Conference on Bryozoa, Vienna 1983. Olsen & Olsen.
26. Wörheide, G., J. N. Hooper, and B. M. Degnan. 2002. Phylogeography of western Pacific *Leucetta 'chagosensis'* (Porifera: Calcarea) from ribosomal DNA sequences: implications for population history and conservation of the Great Barrier Reef World Heritage Area (Australia). Mol Ecol 11:1753-1768.
27. Johnson, M. F. 1978. Recruitment, growth, mortality and seasonal variations in the calcareous sponges *Clathrina coriacea* (Montagu) and *C. blanca* (Miklucho-Maclay) from Santa Catalina island, California. Biologie des spongiaires CNRS Colloque International, Paris. CNRS editorial, 325-334 pp.
28. Johnson, M. F. 1978. Studies on the reproductive cycles of the calcareous sponges *Clathrina coriacea* and *C. blanca*. Marine Biology 50:73-79.
29. Calazans, V. P. S. B., and E. Lanna. 2019. Influence of endogenous and exogenous factors on the reproductive output of a cryptogenic calcareous sponge. Marine Biodiversity 49:2837-2850.
30. Adey, W. H., and J. M. Vassar. 1975. Colonization, succession and growth rates of tropical crustose coralline algae (Rhodophyta, Cryptonemiales). Phycologia 14:55-69.
31. Matsuda, S. 1989. Succession and growth rates of encrusting crustose coralline algae (Rhodophyta, Cryptonemiales) in the upper fore-reef environment off Ishigaki Island, Ryukyu Islands. Coral Reefs 7:185-195.
32. McCoy, S. J., and N. A. Kamenos. 2015. Coralline algae (Rhodophyta) in a changing world: integrating ecological, physiological, and geochemical responses to global change. Journal of Phycology 51:6-24.
33. Caragnano, A., D. Basso, D. E. Jacob, D. Storz, G. Rodondi, F. Benzoni, and E. Dutrieux. 2014. The coralline red alga *Lithophyllum kotschyanum* f. *affine* as proxy of climate variability in the Yemen coast, Gulf of Aden (NW Indian Ocean). Geochimica et Cosmochimica Acta 124:1-17.
34. Heimberg, B. F., and J. E. Winston. 1986. Bryozoans from Bali, Lombok, and Komodo. American Museum novitates ; no. 2847.
35. Chadwick-Furman, N. E., and M. Spiegel. 2000. Abundance and clonal replication in the tropical corallimorpharian *Rhodactis rhodostoma*. Invertebrate Biology 119:351-360.
36. Chadwick-Furman, N. E., M. Spiegel, and I. Nir. 2000. Sexual reproduction in the tropical corallimorpharian *Rhodactis rhodostoma*. Invertebrate Biology 119:361-369.
37. Kuguru, B., N. E. Chadwick, Y. Achituv, K. Zandbank, and D. Tchernov. 2008. Mechanisms of habitat segregation between corallimorpharians: photosynthetic parameters and *Symbiodinium* types. Marine Ecology Progress Series 369:115-129.
38. Elliott, J., Cook, C.B. 1989. Diel variation in prey capture behavior by the corallimorpharian *Discosoma sanctithomae*: mechanical and chemical activation of feeding. The Biological Bulletin 176:218-228.
39. Ahern, K. S., C. R. Ahern, and J. W. Udy. 2008. In situ field experiment shows *Lyngbya majuscula* (cyanobacterium) growth stimulated by added iron, phosphorus and nitrogen. Harmful Algae 7:389-404.

63. McLean, E., and P. Yoshioka. 2008. Substratum effects on the growth and survivorship of the sponge *Desmapsamma anchorata*. Caribbean Journal of Science 44.

64. Murray, J. 2014. Ecology and palaeoecology of benthic foraminifera.

86. Subhan, B., D. Soedharma, D. Arafat, H. Madduppa, F. Rahmawati, A. Ervinia, A. Bramandito, D. Khaerudi, and A. Ghozali. 2017. Pengaruh cahaya terhadap tingkat kelangsungan hidup dan pertumbuhan karang lunak

*Lobophytum strictum* (Octocorallia: Alcyonacea) hasil transplantasi pada system resirkulasi. Jurnal Teknologi Perikanan dan Kelautan 3:35.

87. De Ruyter van Steveninck, E. D., L. L. Van Mulekom, and A. M. Breeman. 1988. Growth inhibition of *Lobophora variegata* (Lamouroux) Womersley by scleractinian corals. Journal of Experimental Marine Biology and Ecology 115:169-178.
88. Mebrahtu, A., J. H. Bruggemann, and M. B. Anneke. 2005. Seasonal patterns of biomass, growth and reproduction in *Dictyota cervicornis* and *Stoechospermum polypodioides* (Dictyotales, Phaeophyta) on a shallow reef flat in the southern Red Sea (Eritrea). Botanica Marina 48:8-17.
89. Asa, S., T. Yeemin, N. Chaitanawisuti, and S. Kritsanapuntu. 2000. Sexual reproduction of a marine sponge, *Petrosia* sp. from coral communities in the Gulf of Thailand. Proceedings 9th International Coral Reef Symposium 1.
90. Huang, D., H. Ou, D. Wang, J. Chen, and S. Ding. 2015. Sexual reproduction of the potentially cultivable *sponge Mycale phyllophila* (Porifera, Demospongiae). Journal of the Marine Biological Association of the United Kingdom 96:1073-1081.
91. Lewis, J. B. 2006. Biology and Ecology of the Hydrocoral *Millepora* on Coral Reefs. Pages 1-55  Advances in Marine Biology. Academic Press.
92. Hwang, S. J., and J. I. Song. 2009. Sexual reproduction of soft coral, *Scleronephthya gracillimum*, (Alcyonacea: Nephtheidae) based on long-term collection from Jejudo Island, Korea. Galaxea, Journal of Coral Reef Studies 11:155-167.
93. Farrant, P. A. 1987. Population dynamics of the temperate Australian soft coral *Capnella gaboensis*. Marine Biology 96:401-407.
94. Gateño, D., Y. Barki, and B. Rinkevich. 2000. Aquarium maintenance of reef octocorals raised from field collected larvae. Aquarium Sciences and Conservation 2:227-236.
95. Babcock, R., and C. Mundy. 1996. Coral recruitment: Consequences of settlement choice for early growth and survivorship in two scleractinians. Journal of Experimental Marine Biology and Ecology 206:179-201.
96. Silva, J. F., P. B. Gomes, E. C. Santana, J. M. Silva, É. P. Lima, A. M. M. Santos, and C. D. Pérez. 2015. Growth of the tropical zoanthid *Palythoa caribaeorum* (Cnidaria: Anthozoa) on reefs in northeastern Brazil. Anais da Academia Brasileira de Ciências 87:985-996.
97. Fadlallah, Y. H., R. H. Karlson, and K. P. Sebens. 1984. A comparative study of sexual reproduction in three species of Panamanian zoanthids (Coelenterata: Anthozoa). Bulletin of Marine Science 35:80-89.
98. Reimer, A. A. 1971. Feeding behavior in the Hawaiian zoanthids *Palythoa* and *Zoanthus*. Pac Sci 25(4): 512-513.
99. Ryland, J. S. 1997. Reproduction in Zoanthidea (Anthozoa: Hexacorallia). Invertebrate Reproduction & Development 31:177-188.
100. Shiroma, E., and J. Reimer. 2010. Eriko Shiroma, James Davis Reimer. 2010. Investigations into the reproductive patterns, ecology and morphology in the zoanthid genus *Palythoa* (Cnidaria: Anthozoa: Hexacorallia) in Okinawa, Japan. Zoological Studies 49: 182-194. Zoological studies 49:182-194.
101. Ng, C. S. L., and L. M. Chou. 2014. Rearing juvenile 'corals of opportunity' in in situ nurseries – A reef rehabilitation approach for sediment-impacted environments. Marine Biology Research 10:833-838.
102. Whalan, S., M. S. Johnson, E. Harvey, and C. Battershill. 2005. Mode of reproduction, recruitment, and genetic subdivision in the brooding sponge *Haliclona* sp. Marine Biology 146:425-433.
103. Rocha, R. J. M., R. Calado, P. Cartaxana, J. Furtado, and J. Serôdio. 2013. Photobiology and growth of leather coral *Sarcophyton* cf. *glaucum* fragments stocked under low light in a recirculated system. Aquaculture 414-415:235-242.
104. Fabricius, K. E. 1995. Slow population turnover in the soft coral genera *Sinularia* and *Sarcophyton* on mid- and outer-shelf reefs of the Great Barrier Reef. Marine Ecology Progress Series 126:145-152.
105. Hellström, M., K. D. Kavanagh, and J. A. H. Benzie. 2010. Multiple spawning events and sexual reproduction in the octocoral *Sarcophyton elegans* (Cnidaria: Alcyonacea) on Lizard Island, Great Barrier Reef. Marine Biology 157:383-392.
106. Benayahu, Y., and Y. Loya. 1986. Sexual reproduction of a soft coral: synchronous and brief annual spawning of *Sarcophyton glaucum* (Quoy & Gaimard, 1933). The Biological Bulletin 170:32-42.
107. Khalesi, M. K., H. H. Beeftink, and R. H. Wijffels. 2011. Energy budget for the cultured, zooxanthellate octocoral *Sinularia flexibilis*. Marine Biotechnology 13:1092-1098.

108. Slattery, M., G. A. Hines, J. Starmer, and V. J. Paul. 1999. Chemical signals in gametogenesis, spawning, and larval settlement and defense of the soft coral *Sinularia polydactyla*. Coral Reefs 18:75-84.
109. Bastidas, C., K. E. Fabricius, and B. L. Willis. 2004. Demographic aspects of the soft coral *Sinularia flexibilis* leading to local dominance on coral reefs. Pages 433-441 *in* Coelenterate Biology 2003. Springer Netherlands, Dordrecht.
110. Mohamed, T., M. M. El-Komi, F. M. Shoukr, and M. A. H. El-Arab. 2017. Growth Rate Evaluation of the Alcyonacean Soft Coral *Sinularia polydactyla* (Ehrenberg, 1834) at Hurghada, Northern Red Sea, Egypt. Journal of Biology, Agriculture and Healthcare 7:42-50.
111. Kahng, S. E., Y. Benayahu, D. Wagner, and N. Rothe. 2008. Sexual reproduction in the invasive octocoral *Carijoa Riisei* in Hawaii. Bulletin of Marine Science 82:1-17.
112. Wagner, D., D. G. Luck, and R. J. Toonen. 2012. Chapter Two - The Biology and Ecology of Black Corals (Cnidaria: Anthozoa: Hexacorallia: Antipatharia). Pages 67-132 *in* M. Lesser, editor. Advances in Marine Biology. Academic Press.
113. Bo, M., C. Di Camillo, A. Addamo, L. Valisano, and G. Bavestrello. 2009. Growth strategies of whip black corals (Cnidaria: Antipatharia) in the Bunaken Marine Park (Celebes Sea, Indonesia). Marine Biodiversity Records 2.
114. Brazeau, D. A., and H. R. Lasker. 1992. Growth rates and growth strategy in a clonal marine invertebrate, the Caribbean octocoral *Briareum asbestinum*. The Biological Bulletin 183:269-277.
115. Gaino, E., and F. Scoccia. 2009. Release of sperm clusters in spheres by the black coral *Cupressopathes pumila* (Anthozoa, Antipatharia). Coral Reefs 28:851-857.
116. Sirison, N., and N. P. Burnett. 2020. *Turbinaria ornata* (Phaeophyceae) varies size and strength to maintain environmental safety factor across flow regimes. Journal of Phycology 56:233-237.
117. Michael, D. G., M. G. Gwendoline, M. Liam, R. Fabio, M. Salvador Valenzuela, C. M. Arthur, C. P. Bruce, L. Anders, M. J. David, B. Ignacio, F. C. Christopher, K. Pier, and J. G. David. 2014. AlgaeBase: An On-line Resource for Algae. Cryptogamie, Algologie 35:105-115.
118. Connell, S. D., M. S. Foster, and L. Airoldi. 2014. What are algal turfs? Towards a better description of turfs. Marine Ecology Progress Series 495:299-307.
119. Benayahu, Y. 1991. Reproduction and developmental pathways of Red Sea Xeniidae (Octocorallia, Alcyonacea). Hydrobiologia 216:125-130.
120. Benayahu, Y., and Y. Loya. 1984. Life history studies on the Red Sea soft coral *Xenia macrospiculata* Gohar, 1940. Annual dynamics of gonadal development. The Biological Bulletin 166:32-43.
121. Ben-David-Zaslow, R., G. Henning, D. K. Hofmann, and Y. Benayahu. 1999. Reproduction in the Red Sea soft coral *Heteroxenia fuscescens*: seasonality and long-term record (1991 to 1997). Marine Biology 133:553-559.
122. Zohar, P., B. Ami, A. Avigdor, and A. Yair. 2004. Initiation of symbiosis between the soft coral *Heteroxenia fuscescens* and its zooxanthellae. Marine Ecology Progress Series 279:113-116.

**Supplementary table 3**. Details on the different fish species identified (highest taxonomic resolution data), their family (major category) and the different trait categories. Fish species with the same FE (Functional entity number) are grouped into the same functional entity. Size (1: 0-7 cm, 2: 7.1-15 cm, 3: 15.1-30 cm, 4: 30.1-50 cm, 5: 50.1-80 cm, 6: > 80 cm). Mob: mobility (1: highly site-attached species (i.e. territorial species), 2: mobile species with a small home range, 3: mobile species with a large home range, 4: widely mobile species with a very large home range). Act: feeding activity (0: diurnal, 1: nocturnal). Greg: gregariousness (1: solitary, 2: pairing, 3: < 20 individuals, 4: >20 individuals. VP: Vertical position (1: benthic, 2: bentho-pelagic, 3: pelagic).

| **Species** | **Family** | **FE** | **Size** | **Mob.** | **Act.** | **Greg.** | **VP** | **Diet** |
|---|---|---|---|---|---|---|---|---|
| ***Chromis_retrofasciata*** | Pomacentridae | #1 | 1 | 2 | 0 | 1 | 1 | Planktivore |
| ***Bodianus_dictynna*** | Labridae | #10 | 2 | 2 | 0 | 1 | 1 | Micro-invertivore |
| ***Chaetodon_ulietensis*** | Chaetodontidae | #10 | 2 | 2 | 0 | 1 | 1 | Micro-invertivore |
| ***Haliocheres_prosopeion*** | Labridae | #10 | 2 | 2 | 0 | 1 | 1 | Micro-invertivore |
| ***Pseudocheilinus_octotaenia*** | Labridae | #10 | 2 | 2 | 0 | 1 | 1 | Micro-invertivore |
| ***Amblyglyphidodon_leucogaster*** | Pomacentridae | #11 | 2 | 2 | 0 | 1 | 1 | Omnivore |
| ***Neopomacentrus_bankieri*** | Pomacentridae | #11 | 2 | 2 | 0 | 1 | 1 | Omnivore |
| ***Chaetodon_lunulatus*** | Chaetodontidae | #12 | 2 | 2 | 0 | 2 | 1 | Corallivore |
| ***Chaetodon_citrinellus*** | Chaetodontidae | #13 | 2 | 2 | 0 | 2 | 1 | Micro-invertivore |
| ***Chaetodon_kleinii*** | Chaetodontidae | #13 | 2 | 2 | 0 | 2 | 1 | Micro-invertivore |
| ***Chaetodon_punctatofasciatus*** | Chaetodontidae | #13 | 2 | 2 | 0 | 2 | 1 | Micro-invertivore |
| ***Ptereleotris_evides*** | Microdesmidae | #14 | 2 | 2 | 0 | 2 | 1 | Planktivore |
| ***Amblyglyphidodon_curacao*** | Pomacentridae | #15 | 2 | 2 | 0 | 3 | 1 | Omnivore |
| ***Centropyge_vrolikii*** | Pomacanthidae | #15 | 2 | 2 | 0 | 3 | 1 | Omnivore |
| ***Pomacentrus_amboinensis*** | Pomacentridae | #15 | 2 | 2 | 0 | 3 | 1 | Omnivore |
| ***Pomacentrus_brachialis*** | Pomacentridae | #15 | 2 | 2 | 0 | 3 | 1 | Omnivore |
| ***Pomacentrus_moluccensis*** | Pomacentridae | #15 | 2 | 2 | 0 | 3 | 1 | Omnivore |
| ***Pomacentrus_nigromarginatus*** | Pomacentridae | #15 | 2 | 2 | 0 | 3 | 1 | Omnivore |
| ***Chromis_atripes*** | Pomacentridae | #16 | 2 | 2 | 0 | 3 | 1 | Planktivore |
| ***Chromis_caudalis*** | Pomacentridae | #16 | 2 | 2 | 0 | 3 | 1 | Planktivore |
| ***Chromis_ternatensis*** | Pomacentridae | #16 | 2 | 2 | 0 | 3 | 1 | Planktivore |
| ***Amblyglyphidodon_aureus*** | Pomacentridae | #17 | 2 | 2 | 0 | 3 | 2 | Planktivore |
| ***Chromis_cinerascens*** | Pomacentridae | #17 | 2 | 2 | 0 | 3 | 2 | Planktivore |
| ***Chromis_margaritifer*** | Pomacentridae | #17 | 2 | 2 | 0 | 3 | 2 | Planktivore |
| ***Chromis_weberi*** | Pomacentridae | #17 | 2 | 2 | 0 | 3 | 2 | Planktivore |
| ***Neopomacentrus_anabatoides*** | Pomacentridae | #18 | 2 | 2 | 0 | 4 | 1 | Omnivore |
| ***Neopomacentrus_cyanomos*** | Pomacentridae | #18 | 2 | 2 | 0 | 4 | 1 | Omnivore |
| ***Acanthochromis_polyacanthus*** | Pomacentridae | #19 | 2 | 2 | 0 | 4 | 2 | Planktivore |

| | | | | | | | | |
|---|---|---|---|---|---|---|---|---|
| ***Chromis_amboinensis*** | Pomacentridae | #19 | 2 | 2 | 0 | 4 | 2 | Planktivore |
| ***Chromis_atripectoralis*** | Pomacentridae | #19 | 2 | 2 | 0 | 4 | 2 | Planktivore |
| ***Canthigaster_valentini*** | Tetraodontidae | #2 | 2 | 1 | 0 | 1 | 1 | Micro-invertivore |
| ***Labroides_bicolor*** | Labridae | #2 | 2 | 1 | 0 | 1 | 1 | Micro-invertivore |
| ***Labroides_pectoralis*** | Labridae | #2 | 2 | 1 | 0 | 1 | 1 | Micro-invertivore |
| ***Spratelloides_gracillis*** | Clupeidae | #20 | 2 | 3 | 0 | 4 | 3 | Planktivore |
| ***Dischistodus_melanotus*** | Labridae | #21 | 3 | 1 | 0 | 1 | 1 | Grazer/Detritivore |
| ***Balistapus_undulatus*** | Balistidae | #22 | 3 | 1 | 0 | 1 | 1 | Macro-invertivore |
| ***Paracirrhites_forsteri*** | Cirrhitidae | #22 | 3 | 1 | 0 | 1 | 1 | Macro-invertivore |
| ***Neoglyphidodon_melas*** | Pomacentridae | #23 | 3 | 1 | 0 | 1 | 1 | Omnivore |
| ***Chaetodon_baronessa*** | Chaetodontidae | #24 | 3 | 1 | 0 | 2 | 1 | Corallivore |
| ***Amphiprion_sebae*** | Pomacentridae | #25 | 3 | 1 | 0 | 2 | 1 | Omnivore |
| ***Synodus_dermatogenys*** | Synodontidae | #26 | 3 | 1 | 1 | 1 | 1 | Piscivore |
| ***Amanses_scopas*** | Monacanthidae | #27 | 3 | 2 | 0 | 1 | 1 | Corallivore |
| ***Chaetodon_trifascialis*** | Chaetodontidae | #27 | 3 | 2 | 0 | 1 | 1 | Corallivore |
| ***Labrichthys_unilineatus*** | Labridae | #27 | 3 | 2 | 0 | 1 | 1 | Corallivore |
| ***Acanthurus_japonicus*** | Acanthuridae | #28 | 3 | 2 | 0 | 1 | 1 | Grazer/Detritivore |
| ***Acanthurus_nigricans*** | Acanthuridae | #28 | 3 | 2 | 0 | 1 | 1 | Grazer/Detritivore |
| ***Bodianus_axillaris*** | Labridae | #29 | 3 | 2 | 0 | 1 | 1 | Micro-invertivore |
| ***Bodianus_mesothorax*** | Labridae | #29 | 3 | 2 | 0 | 1 | 1 | Micro-invertivore |
| ***Chaetodon_speculum*** | Chaetodontidae | #29 | 3 | 2 | 0 | 1 | 1 | Micro-invertivore |
| ***Choerodon_jordani*** | Labridae | #29 | 3 | 2 | 0 | 1 | 1 | Micro-invertivore |
| ***Coris_caudimacula*** | Labridae | #29 | 3 | 2 | 0 | 1 | 1 | Micro-invertivore |
| ***Haliocheres_melanochir*** | Labridae | #29 | 3 | 2 | 0 | 1 | 1 | Micro-invertivore |
| ***Haliocheres_podostigma*** | Labridae | #29 | 3 | 2 | 0 | 1 | 1 | Micro-invertivore |
| ***Pomacanthus_navarchus*** | Pomacanthidae | #29 | 3 | 2 | 0 | 1 | 1 | Micro-invertivore |
| ***Pygoplites_diacanthus*** | Pomacanthidae | #29 | 3 | 2 | 0 | 1 | 1 | Micro-invertivore |
| ***Sufflamen_chrysopterum*** | Balistidae | #29 | 3 | 2 | 0 | 1 | 1 | Micro-invertivore |
| ***Neoglyphidodon_nigroris*** | Pomacentridae | #3 | 2 | 1 | 0 | 1 | 1 | Omnivore |
| ***Thalassoma_hardwicke*** | Labridae | #30 | 3 | 2 | 0 | 1 | 2 | Micro-invertivore |
| ***Chaetodon_melanotus*** | Chaetodontidae | #31 | 3 | 2 | 0 | 2 | 1 | Corallivore |
| ***Chaetodon_ornatissimus*** | Chaetodontidae | #31 | 3 | 2 | 0 | 2 | 1 | Corallivore |
| ***Siganus_vulpinus*** | Siganidae | #32 | 3 | 2 | 0 | 2 | 1 | Grazer/Detritivore |
| ***Chaetodon_auriga*** | Chaetodontidae | #33 | 3 | 2 | 0 | 2 | 1 | Micro-invertivore |
| ***Chaetodon_epphiphium*** | Chaetodontidae | #33 | 3 | 2 | 0 | 2 | 1 | Micro-invertivore |
| ***Chaetodon_rafflesi*** | Chaetodontidae | #33 | 3 | 2 | 0 | 2 | 1 | Micro-invertivore |
| ***Heniochus_chrysostomus*** | Chaetodontidae | #33 | 3 | 2 | 0 | 2 | 1 | Micro-invertivore |

| | | | | | | | | |
|---|---|---|---|---|---|---|---|---|
| ***Heniochus_varius*** | Chaetodontidae | #33 | 3 | 2 | 0 | 2 | 1 | Micro-invertivore |
| ***Chaetodon_vagabundus*** | Chaetodontidae | #34 | 3 | 2 | 0 | 2 | 1 | Omnivore |
| ***Zanclus_cornutus*** | Zanclidae | #35 | 3 | 2 | 0 | 2 | 2 | Micro-invertivore |
| ***Chaetodon_lunula*** | Chaetodontidae | #36 | 3 | 2 | 0 | 3 | 1 | Micro-invertivore |
| ***Chaetodon_unimaculatus*** | Chaetodontidae | #36 | 3 | 2 | 0 | 3 | 1 | Micro-invertivore |
| ***Chaetodontoplus_mesoleucus*** | Chaetodontidae | #36 | 3 | 2 | 0 | 3 | 1 | Micro-invertivore |
| ***Chromis_notata*** | Pomacentridae | #37 | 3 | 2 | 0 | 3 | 1 | Planktivore |
| ***Forcipiger_flavissimus*** | Chaetodontidae | #38 | 3 | 2 | 0 | 3 | 2 | Micro-invertivore |
| ***Thalassoma_amblycephalum*** | Labridae | #38 | 3 | 2 | 0 | 3 | 2 | Micro-invertivore |
| ***Thalassoma_jansenii*** | Labridae | #38 | 3 | 2 | 0 | 3 | 2 | Micro-invertivore |
| ***Chromis_analis*** | Pomacentridae | #39 | 3 | 2 | 0 | 3 | 2 | Planktivore |
| ***Chromis_xanthura*** | Pomacentridae | #39 | 3 | 2 | 0 | 3 | 2 | Planktivore |
| ***Labroides_dimidiatus*** | Labridae | #4 | 2 | 1 | 0 | 2 | 1 | Micro-invertivore |
| ***Abudefduf_vaigensis*** | Pomacentridae | #40 | 3 | 2 | 0 | 4 | 2 | Planktivore |
| ***Hemitaurichthys_polylepis*** | Chaetodontidae | #40 | 3 | 2 | 0 | 4 | 2 | Planktivore |
| ***Myripristis_botche*** | Holocentridae | #41 | 3 | 2 | 1 | 2 | 2 | Planktivore |
| ***Sargocentron_diadema*** | Holocentridae | #42 | 3 | 2 | 1 | 3 | 1 | Macro-invertivore |
| ***Scolopsis_affinis*** | Nemipteridae | #43 | 3 | 2 | 1 | 3 | 1 | Micro-invertivore |
| ***Scolopsis_bilineatus*** | Nemipteridae | #43 | 3 | 2 | 1 | 3 | 1 | Micro-invertivore |
| ***Scolopsis_lineatus*** | Nemipteridae | #43 | 3 | 2 | 1 | 3 | 1 | Micro-invertivore |
| ***Pempheris_oualiensis*** | Pempheridae | #44 | 3 | 2 | 1 | 3 | 1 | Omnivore |
| ***Sargocentron_microstoma*** | Holocentridae | #45 | 3 | 2 | 1 | 3 | 1 | Piscivore |
| ***Myripristis_amaena*** | Holocentridae | #46 | 3 | 2 | 1 | 3 | 2 | Planktivore |
| ***Scarus_tricolor*** | Scaridae | #47 | 3 | 3 | 0 | 1 | 1 | Excavator/scraper |
| ***Acanthurus_pyroferus*** | Acanthuridae | #48 | 3 | 3 | 0 | 1 | 1 | Grazer/Detritivore |
| ***Ctenochaetus_striatus*** | Acanthuridae | #48 | 3 | 3 | 0 | 1 | 1 | Grazer/Detritivore |
| ***Ostracion_meleagris*** | Ostraciidae | #49 | 3 | 3 | 0 | 1 | 1 | Micro-invertivore |
| ***Amphiprion_clarkii*** | Pomacentridae | #5 | 2 | 1 | 0 | 2 | 1 | Omnivore |
| ***Amphiprion_frenatus*** | Pomacentridae | #5 | 2 | 1 | 0 | 2 | 1 | Omnivore |
| ***Amphiprion_melanopus*** | Pomacentridae | #5 | 2 | 1 | 0 | 2 | 1 | Omnivore |
| ***Oxycheilinus_diagramma*** | Labridae | #50 | 3 | 3 | 0 | 1 | 1 | Piscivore |
| ***Chlorurus_japanensis*** | Scaridae | #51 | 3 | 3 | 0 | 3 | 1 | Excavator/scraper |
| ***Acanthurus_nigrofuscus*** | Acanthuridae | #52 | 3 | 3 | 0 | 3 | 1 | Grazer/Detritivore |
| ***Zebrasoma_rostratum*** | Acanthuridae | #52 | 3 | 3 | 0 | 3 | 1 | Grazer/Detritivore |
| ***Acanthurus_thompsoni*** | Acanthuridae | #53 | 3 | 3 | 0 | 3 | 2 | Planktivore |
| ***Pterocaesio_pisang*** | Caesionidae | #54 | 3 | 3 | 0 | 4 | 2 | Planktivore |
| ***Pterocaesio_tessellata*** | Caesionidae | #54 | 3 | 3 | 0 | 4 | 2 | Planktivore |
| ***Pterocaesio_tile*** | Caesionidae | #54 | 3 | 3 | 0 | 4 | 2 | Planktivore |

| | | | | | | | | |
|---|---|---|---|---|---|---|---|---|
| ***Gnathodentex_aurolineatus*** | Lethrinidae | #55 | 3 | 3 | 1 | 4 | 2 | Piscivore |
| ***Arothron_hispidus*** | Tetraodontidae | #56 | 4 | 2 | 0 | 1 | 1 | Macro-invertivore |
| ***Anampses_geographicus*** | Labridae | #57 | 4 | 2 | 0 | 1 | 1 | Micro-invertivore |
| ***Sufflamen_fraenatus*** | Balistidae | #57 | 4 | 2 | 0 | 1 | 1 | Micro-invertivore |
| ***Thalassoma_lunare*** | Labridae | #58 | 4 | 2 | 0 | 1 | 2 | Micro-invertivore |
| ***Hologymnosus_doliatus*** | Labridae | #59 | 4 | 2 | 0 | 1 | 2 | Piscivore |
| ***Dascyllus_aruanus*** | Pomacentridae | #6 | 2 | 1 | 0 | 3 | 1 | Omnivore |
| ***Dascyllus_reticulatus*** | Pomacentridae | #6 | 2 | 1 | 0 | 3 | 1 | Omnivore |
| ***Dascyllus_trimaculatus*** | Pomacentridae | #6 | 2 | 1 | 0 | 3 | 1 | Omnivore |
| ***Myripristis_adusta*** | Holocentridae | #60 | 4 | 2 | 1 | 1 | 2 | Planktivore |
| ***Sargocentron_caudimaculatum*** | Holocentridae | #61 | 4 | 2 | 1 | 3 | 1 | Piscivore |
| ***Sargocentron_praslin*** | Holocentridae | #61 | 4 | 2 | 1 | 3 | 1 | Piscivore |
| ***Sargocentron_rubrum*** | Holocentridae | #61 | 4 | 2 | 1 | 3 | 1 | Piscivore |
| ***Zebrasoma_velifera*** | Acanthuridae | #62 | 4 | 3 | 0 | 1 | 1 | Browser |
| ***Scarus_dimidiatus*** | Scaridae | #63 | 4 | 3 | 0 | 1 | 1 | Excavator/scraper |
| ***Scarus_ferrugines*** | Scaridae | #63 | 4 | 3 | 0 | 1 | 1 | Excavator/scraper |
| ***Scarus_frenatus*** | Scaridae | #63 | 4 | 3 | 0 | 1 | 1 | Excavator/scraper |
| ***Scarus_niger*** | Scaridae | #63 | 4 | 3 | 0 | 1 | 1 | Excavator/scraper |
| ***Scarus_oviceps*** | Scaridae | #63 | 4 | 3 | 0 | 1 | 1 | Excavator/scraper |
| ***Scarus_scaber*** | Scaridae | #63 | 4 | 3 | 0 | 1 | 1 | Excavator/scraper |
| ***Scarus_viridifucatus*** | Scaridae | #63 | 4 | 3 | 0 | 1 | 1 | Excavator/scraper |
| ***Cheilinus_fasciatus*** | Labridae | #64 | 4 | 3 | 0 | 1 | 1 | Macro-invertivore |
| ***Hemigymnus_melapterus*** | Labridae | #64 | 4 | 3 | 0 | 1 | 1 | Macro-invertivore |
| ***Parupeneus_indicus*** | Mullidae | #64 | 4 | 3 | 0 | 1 | 1 | Macro-invertivore |
| ***Parupeneus_multifasciatus*** | Mullidae | #64 | 4 | 3 | 0 | 1 | 1 | Macro-invertivore |
| ***Parupeneus_pleurostigma*** | Mullidae | #64 | 4 | 3 | 0 | 1 | 1 | Macro-invertivore |
| ***Anampses_caeruleopunctatus*** | Labridae | #65 | 4 | 3 | 0 | 1 | 1 | Micro-invertivore |
| ***Cantherhines_dumerilii*** | Monacanthidae | #65 | 4 | 3 | 0 | 1 | 1 | Micro-invertivore |
| ***Hologymnosus_annulatus*** | Labridae | #66 | 4 | 3 | 0 | 1 | 1 | Piscivore |
| ***Balistoides_conspicillum*** | Balistidae | #67 | 4 | 3 | 0 | 1 | 2 | Macro-invertivore |
| ***Melchthys_vidua*** | Balistidae | #68 | 4 | 3 | 0 | 1 | 2 | Planktivore |
| ***Chlorurus_capistratoides*** | Scaridae | #69 | 4 | 3 | 0 | 3 | 1 | Excavator/scraper |
| ***Chlorurus_sordidus*** | Scaridae | #69 | 4 | 3 | 0 | 3 | 1 | Excavator/scraper |
| ***Chlorurus_spilurus*** | Scaridae | #69 | 4 | 3 | 0 | 3 | 1 | Excavator/scraper |
| ***Pseudanthias_squamipinnis*** | Serranidae | #7 | 2 | 1 | 0 | 3 | 2 | Planktivore |
| ***Pseudoanthias_cooperi*** | Serranidae | #7 | 2 | 1 | 0 | 3 | 2 | Planktivore |
| ***Acanthurus_blochii*** | Acanthuridae | #70 | 4 | 3 | 0 | 3 | 1 | Grazer/Detritivore |
| ***Zebrasoma_scopas*** | Acanthuridae | #70 | 4 | 3 | 0 | 3 | 1 | Grazer/Detritivore |

| | | | | | | | | |
|---|---|---|---|---|---|---|---|---|
| ***Lethrinus_harak*** | Lethrinidae | #71 | 4 | 3 | 0 | 3 | 1 | Piscivore |
| ***Naso_lituratus*** | Acanthuridae | #72 | 4 | 3 | 0 | 3 | 2 | Browser |
| ***Naso_sp*** | Acanthuridae | #72 | 4 | 3 | 0 | 3 | 2 | Browser |
| ***Odonus_niger*** | Balistidae | #73 | 4 | 3 | 0 | 3 | 2 | Planktivore |
| ***Parupeneus_bifasciatus*** | Mullidae | #74 | 4 | 3 | 1 | 1 | 1 | Macro-invertivore |
| ***Lutjanus_fulvus*** | Lutjanidae | #75 | 4 | 3 | 1 | 1 | 2 | Piscivore |
| ***Lutjanus_carponotatus*** | Lutjanidae | #76 | 4 | 3 | 1 | 3 | 2 | Piscivore |
| ***Lutjanus_lutjanus*** | Lutjanidae | #77 | 4 | 3 | 1 | 4 | 1 | Piscivore |
| ***Caesio_caerulaurea*** | Caesionidae | #78 | 4 | 4 | 0 | 4 | 3 | Planktivore |
| ***Caesio_lunaris*** | Caesionidae | #78 | 4 | 4 | 0 | 4 | 3 | Planktivore |
| ***Gymnothorax_eurostus*** | Muraenidae | #79 | 5 | 1 | 1 | 1 | 1 | Piscivore |
| ***Pseudoanthias_dispar*** | Serranidae | #8 | 2 | 1 | 0 | 4 | 2 | Planktivore |
| ***Aulostomus_chinensis*** | Aulostomidae | #80 | 5 | 2 | 0 | 1 | 1 | Piscivore |
| ***Balistoides_viridescens*** | Balistidae | #81 | 5 | 2 | 0 | 1 | 2 | Macro-invertivore |
| ***Cephalopholis_argus*** | Serranidae | #82 | 5 | 2 | 0 | 3 | 1 | Piscivore |
| ***Myripristis_murdjan*** | Holocentridae | #83 | 5 | 2 | 1 | 2 | 2 | Planktivore |
| ***Scarus_rubroviolaceous*** | Scaridae | #84 | 5 | 3 | 0 | 1 | 1 | Excavator/scraper |
| ***Parupeneus_barberinus*** | Mullidae | #85 | 5 | 3 | 0 | 1 | 1 | Macro-invertivore |
| ***Bodianus_perditio*** | Labridae | #86 | 5 | 3 | 0 | 1 | 1 | Micro-invertivore |
| ***Naso_vlamingi*** | Acanthuridae | #87 | 5 | 3 | 0 | 1 | 2 | Omnivore |
| ***Chlorurus_strongylocephalus*** | Scaridae | #88 | 5 | 3 | 0 | 3 | 1 | Excavator/scraper |
| ***Lethrinus_obsoletus*** | Lethrinidae | #89 | 5 | 3 | 0 | 3 | 1 | Piscivore |
| ***Diproctacanthus_xanthurus*** | Labridae | #9 | 2 | 2 | 0 | 1 | 1 | Corallivore |
| ***Lutjanusmonostigma*** | Lutjanidae | #90 | 5 | 3 | 1 | 1 | 1 | Piscivore |
| ***Plectorhincus_lineatus*** | Haemulidae | #91 | 5 | 3 | 1 | 3 | 2 | Micro-invertivore |
| ***Platax_teira*** | Ephippidae | #92 | 5 | 4 | 0 | 3 | 3 | Omnivore |
| ***Caranx_sp*** | Carangidae | #93 | 5 | 4 | 0 | 3 | 3 | Piscivore |
| ***Macolor_macularis*** | Lutjanidae | #94 | 5 | 4 | 1 | 3 | 2 | Piscivore |
| ***Coris_aygula*** | Labridae | #95 | 6 | 3 | 0 | 1 | 1 | Macro-invertivore |
| ***Fistularia_commersonii*** | Fistulariidae | #96 | 6 | 3 | 0 | 1 | 2 | Piscivore |
| ***Sphyraena_qenie*** | Sphyraenidae | #97 | 6 | 4 | 1 | 4 | 3 | Piscivore |
